# Coronavirus Nucleocapsid Protein Enhances the binding of p-PKCα to RACK1: Implications for Inhibition of Nucleocytoplasmic Trafficking and Suppression of the Innate Immune Response

**DOI:** 10.1101/2024.03.06.583677

**Authors:** Wenxiang Xue, Hongyan Chu, Jiehuang Wang, Yingjie Sun, Xusheng Qiu, Cuiping Song, Lei Tan, Chan Ding, Ying Liao

## Abstract

The hallmark of coronavirus infection lies in its ability to evade host immune defenses, a process intricately linked to the nuclear entry of transcription factors crucial for initiating the expression of antiviral genes. Central to this evasion strategy is the manipulation of the nucleocytoplasmic trafficking system, which serves as an effective target for the virus to modulate the expression of immune response-related genes. In this investigation, we discovered that infection with the infectious bronchitis virus (IBV) dynamically impedes the nuclear translocation of several transcription factors such as IRF3, STAT1, STAT2, NF-κB p65, and the p38 mitogen-activated protein kinase (MAPK), leading to compromised transcriptional induction of key antiviral genes such as IFNβ, IFITM3, and IL-8. Further examination revealed that during the infection process, components of the nuclear pore complex (NPC), particularly FG-Nups (such as NUP62, NUP153, NUP42, and TPR), undergo cytosolic dispersion from the nuclear envelope; NUP62 undergoes phosphorylation, and NUP42 exhibits a mobility shift in size. These observations suggest a disruption in nucleocytoplasmic trafficking. Screening efforts identified the IBV nucleocapsid protein (N) as the agent responsible for the cytoplasmic distribution of FG-Nups, subsequently hindering the nuclear entry of transcription factors and suppressing the expression of antiviral genes. Interactome analysis further revealed that the IBV N protein interacts with the scaffold protein RACK1, facilitating the recruitment of activated protein kinase C alpha (p-PKCα) to RACK1 and relocating the RACK1-PKCα complex to the cytoplasm. These observations are conserved across pan-coronaviruses N proteins. Concurrently, the presence of both RACK1 and PKCα/β proved essential for the phosphorylation and cytoplasmic dispersion of NUP62, the suppression of antiviral cytokine expression, and efficient virus replication. These findings unveil a novel, highly effective, and evolutionarily conserved mechanism.

**Author summary:** Coronaviruses employ diverse strategies to suppress the host innate immune defense. In this study, we uncovered a novel and highly effective strategy utilized by pan-coronaviruses to inhibit the innate immune response. Specifically, we found that the coronavirus N protein facilitates the binding of p-PKCα to RACK1, leading to the phosphorylation of NUP62 and the cytoplasmic redistribution of multiple FG-Nups. This phenomenon is accompanied by the disruption of nuclear translocation of several innate immune response-related transcription factors and suppression of antiviral/pro-inflammatory genes expression. Our research represents the first elucidation of how the N protein targets and impairs NPC function through the promotion of RACK1-PKCα interaction and NUP62 phosphorylation/disassembly. This discovery unveils a novel mechanism employed by pan-coronaviruses to counteract the host immune response.

## Introduction

The trafficking of proteins and RNA between the nucleus and cytoplasm through nuclear pore complex (NPC) is pivotal for numerous cellular functions, such as gene transcription, RNA and ribosomal subunits export, protein translation, and antiviral innate immunity [1–4]. Comprising the nuclear envelope (NE), NPC, and nuclear transport receptors, the nucleocytoplasmic trafficking system is intricately structured. The outer nuclear membrane (ONM) is contiguous with the endoplasmic reticulum (ER) and shares similarities with it, while the inner nuclear membrane (INM) faces the nucleoplasm and provides anchoring sites for chromatin and the nuclear lamina [5, 6]. NPC, acting as the exclusive gateway controlling molecules transport into and out of the nucleus, consists of approximately 30 different nucleoporins (Nups) with a total molecular mass ∼110 MDa [7–9]. Structurally, NPC features an eight-fold symmetric central core surrounding a transport channel. Nups extending from the central core into the cytoplasm form cytoplasmic filaments, while nuclear-side filaments interconnect to create the nuclear basket. Within the central transport channel, Nups with repeating sequences rich in phenylalanine (Phe) and glycine (Gly), known as FG-Nups, form a cohesive meshwork acting as a permeable barrier to regulate cargo movement. To cross the nuclear pore, proteins over 40 kDa rely on nuclear transport receptors such as importin α, importin β or transportins. The small Ras-like GTPase Ran, which cycles between GDP-bound and GTP-bound states, regulating the formation and disassembly of nuclear transport receptors with cargo proteins or RNA [10]. The NPC dynamics is directly regulated by cell cycle-dependent phosphorylation [11, 12]. Hyperphosphorylation of the gatekeeper NUP98 and the NUP53 by cyclin-dependent kinase 1 (CDK1) and polo-like kinase 1 (PLK1) is a crucial step promoting NPC disintegration [13, 14]. Several Nups, including NUP153, NUP214, and NUP358, undergo phosphorylated throughout the cell cycle and become hyperphosphorylated during M phase, with CDK1 or other kinases likely playing pivotal roles in this process [15]. Dephosphorylation nuclear envelope proteins by the sequential activated phosphatases is essential for correct NPC and nuclear envelope reassembly. Two main phosphatases, protein phosphatase 1 (PP1) and protein phosphatase 2A (PP2A), are involved in the dephosphorylation of Nups and lamins [16, 17]. However, detailed mechanisms underlying the relationship between Nups phosphorylation and NPC disassembly remain unclear.

The NPC governs the nuclear translocation of key signaling transcription transducers crucial for activating the production of antiviral genes, including IFNs, IFN-stimulated genes (ISGs), and pro-inflammatory cytokines. The type I IFN signaling pathway constitutes a central component of the antiviral innate immune response, with its activation dependent on the nuclear translocation of IFN regulatory factor 3 (IRF3) to induce the expression of IFNα/β. These type I IFNs subsequently activate the Janus kinase/signal transducer and activator of transcription (JAK/STAT) pathway upon binding to cell surface receptors, facilitating the nuclear translocation of STAT1/STAT2 and ultimately stimulating the expression of numerous antiviral ISGs [18, 19]. In response to infection signals, the transcription factor NF-κB (comprising p50 and p65 subunits) undergoes phosphorylation and translocation into nucleus to induce the expression of various pro-inflammatory cytokines [20]. The phosphorylation and nuclear translocation of p38 MAPK, triggered by stress stimuli or infection, contribute to inflammation by phosphorylating several nuclear transcription regulators and regulating the stress related transcription processes [21, 22]. In addition to importing transcription factors into the nucleus, the NPC serves as the gate for exporting mRNA from the nucleus into the cytoplasm, including mRNA encoding antiviral genes. Consequently, the host nucleocytoplasmic trafficking system represents an effective target for viruses to regulate the host antiviral gene expression and suppress the immune response.

Coronaviruses pose a significant global health threat to human beings, leading to substantial economic losses. This RNA virus family possesses a positive-sense, single-stranded genome ranging from 25 to 32 kb. Approximately two-thirds of the 5’ genome encode viral replicase polyproteins 1a and 1ab, which are subsequently cleaved into 15-16 mature nonstructural proteins (nsp) crucial for viral replication by internal proteases. The remaining one-third of the 3’ genome encodes structural proteins, including spike (S), envelope (E), membrane (M), nucleocapsid (N), and accessory proteins [23]. Highly pathogenic coronaviruses such as SARS-CoV, SARS-CoV-2, and MERS-CoV cause severe atypical pneumonia, accompanied by coughing and high fever, often resulting in high mortality rates. Conversely, mild pathogenic coronaviruses HCoV-229E, HCoV-OC43, HCoV-NL63, and HKU1 typically induce common cold symptoms and mild upper respiratory disease [23, 24]. Additionally, coronaviruses are also responsible for various infectious diseases in economic animals. For instance, *gamma-coronavirus* IBV causes highly contagious diseases in chickens, manifesting as bronchitis, nephritis, and fallopian tube injury, leading to a significant decrease in laying rate and chicken production. Since its discovery in the 1930s, IBV has persisted as a significant pathogen, posing a continuous threaten in poultry farms [25]. Despite the extensive vaccination effort, controlling this disease remains challenging due to the ongoing emergence of new serotypes and variants [26, 27]. Since its discovery in 1971 [28], porcine epidemic diarrhea virus (PEDV) has posed a substantial threat as a major pathogen in the pig farming industry, causing clinical symptoms such as vomiting and severe diarrhea, causing clinical symptoms such as vomiting and severe diarrhea. The mortality rate among infected suckling piglet can reach100% [29, 30].

Delayed IFN response is a common observation during coronavirus infection, including PEDV [31], SARS-CoV [32], SARS-CoV-2 [33, 34], MERS-CoV [35], MHV [36], IBV [37] and PDCoV [38]. Several excellent review articles have extensively discussed how various proteins encoded by different genera of coronaviruses antagonize the innate immune system through diverse strategies [18, 39–41]. However, research into targeting the nuclear transport system to suppress innate immunity has been limited to SARS-CoV and SARS-CoV-2. Specifically, SARS-CoV ORF6 interacts with importin α1 (KPNA2) and competes with importin α5 (KPNA1) for binding to importin β1 (KPNB1), sequestering importin α1 and importin β1 at the ER/Golgi membrane and blocking the nuclear translocation of STAT1 [42]. Similarly, SARS-CoV-2 ORF6 inhibits STAT1/2 nuclear translocation by obstructing the interaction of importin β1 with NUP98, a major NUP involved in nuclear transport cycle [43]. ORF6 is an accessory protein unique to SARS-CoV and SARS-CoV-2, absent in other coronaviruses. Whether there are evolutionary conserved coronavirus proteins involved in modulating the nuclear trafficking system warrants further investigation.

In this study, we utilized IBV as the model to explore the shared mechanisms utilized by pan-coronaviruses to disrupt the host nuclear transport system. We observed that IBV infection triggered cytoplasmic dispersion of several FG-Nups and hindered the nuclear ingress of various transcription factors and p38 MAPK, consequently impeding the transcriptional activation of antiviral and pro-inflammatory genes. Further study revealed that the IBV N protein was accountable for the disturbance of nucleocytoplasmic trafficking by enhancing the association of p-PKCα with scaffold protein RACK1, leading to relocation of the p-PKCα-RACK1 complex to cytoplasm. This promoted the phosphorylation and cytoplasmic dispersion of NUP62, ultimately impeding the nuclear import of several transcription factors and dampening the transcription of antiviral/pro-inflammatory genes. Importantly, this novel function was conserved among N proteins from four genera of coronaviruses.

## Results

### IBV infection disrupts the nuclear translocation of transcription factors and antagonizes the expression of antiviral and pro-inflammatory cytokines

The previous report has demonstrated that IBV inhibits IFNβ-mediated nuclear translocation of STAT1 at the late stages of infection [44]. In this study, we explored whether IBV also impedes the nuclear translocation of additional transcription factors, including IRF3, STAT1, STAT2, and p65. Initially, Vero cells, an IBV Beaudette strain adapted cell line, were infected with 1 MOI of IBV, followed by poly(I:C) transfection to induce IRF3 nuclear translocation. Immunofluorescence analysis depicted in Fig 1A (left panel) revealed that throughout the infection process (6-18 h.p.i.), IBV infection failed to induce the nuclear entry of IRF3; unlike poly(I:C) transfection which effectively stimulated IRF3 nuclear translocation. Intriguingly, in IBV-infected cells, IRF3 was predominantly distributed in the cytoplasm, indicating that IBV infection hinders the poly(I:C) stimulated IRF3 nuclear translocation (Fig 1A, left panel). Since Vero cells lack IFN, they are suitable for the study of JAK-STAT pathway [45, 46]. Immunofluorescence analysis demonstrated that IBV infection did not trigger the nuclear translocation of both STAT1 and STAT2, whereas IFNβ successfully induced their nuclear translocation. In IBV-infected Vero cells, the nuclear translocation of STAT1 and STAT2 stimulated by IFNβ was impeded, indicated by the diffuse signals of both transcription factors observed in both the cytoplasm and the nucleus (Fig 1A, middle panel). The inhibition of nuclear translocation of aforementioned transcription factors by IBV infection led us to investigate whether the virus also inhibits the nuclear translocation of additional transcription factors or transcription transducers. Consequently, the nuclear translocation of pro-inflammatory transcription factor NF-κB subunit p65 and p38 MAPK was examined. As illustrated in Fig 1A (right panel), IBV infection did not promote the nuclear translocation of p65 and p38 MAPK at 6 h.p.i. and 12 h.p.i.; however, along with the infection progressed, an increasing amount of p65 and p38 MAPK entered the nucleus at 18 h.p.i.. These findings suggest that the virus suppresses inflammatory transcription events during the early stages of infection to facilitate successful infection; however, inflammation is ultimately induced at the late stage of infection. Both TNFα treatment and UV irradiation effectively promoted the nuclear entry of p65 and p38 MAPK, whereas IBV infection led to a proportion of p65 and p38 MAPK remaining in the cytoplasm, despite the detection of intense nuclear signals. The inhibition of nuclear translocation of IRF3, STAT2, and p65 by IBV infection was further validated in host chicken embryo fibroblast DF-1 (S1A-B Fig). Concurrently, the transcription of IFNβ (induced by IRF3), IFITM3 (an ISG induced by STAT1/2), and pro-inflammatory cytokine IL-8 (induced by NF-κB and p38 MAPK) was suppressed by IBV infection in DF-1, which possesses a complete IFN signaling pathway (Fig 1B). Collectively, these results demonstrate that IBV infection suppresses nuclear entry of multiple transcription factors, particularly those involved in the antiviral IFN pathways.

**Fig 1.**
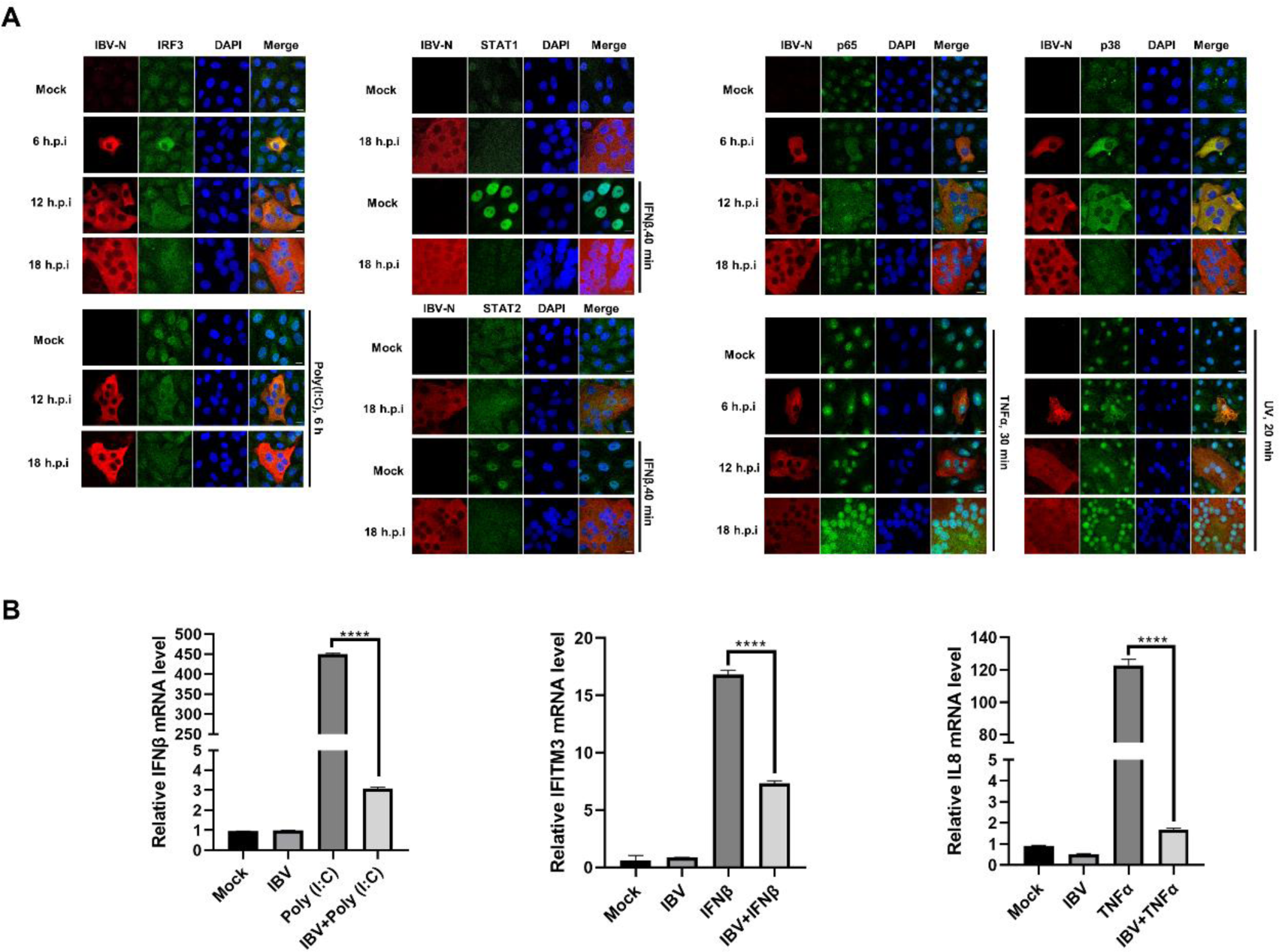
IBV infection impairs nuclear translocation of transcription factors and suppresses transcription of antiviral genes and pro-inflammatory genes. (A) Vero cells were infected with IBV at an MOI of 1, followed by transfection with poly(I:C) (20 μg/mL) for 6 h, treatment with IFNβ (1000 IU/mL) for 40 min, treatment with TNFα (20 ng/mL) for 30 min, or exposure to UV irradiation (1.92 J/cm2) for 20 min. Mock-infected cells served as the control group. Cells were harvested at the indicated time points and subjected to immunofluorescence analysis. Representative images from three independent experiments are shown. Scale bars: 10 μm. (B) DF-1 cells were infected with IBV at an MOI of 5 for 2 h, followed by transfection with poly(I:C) (20 μg/mL) or treatment with TNFα (20 ng/mL) for 6 h. Cells were harvested at 8 h post-infection (h.p.i.) and subjected to qRT-PCR analysis. For detection of IFITM3 expression, DF-1 cells were infected with IBV at an MOI of 5, followed by treatment with IFNβ (1000 IU/mL) for 6 h at 6 h.p.i. Cells were harvested at 12 h.p.i. and subjected to qRT-PCR analysis. Mock-infected cells served as the control group.

**S1 Fig.**
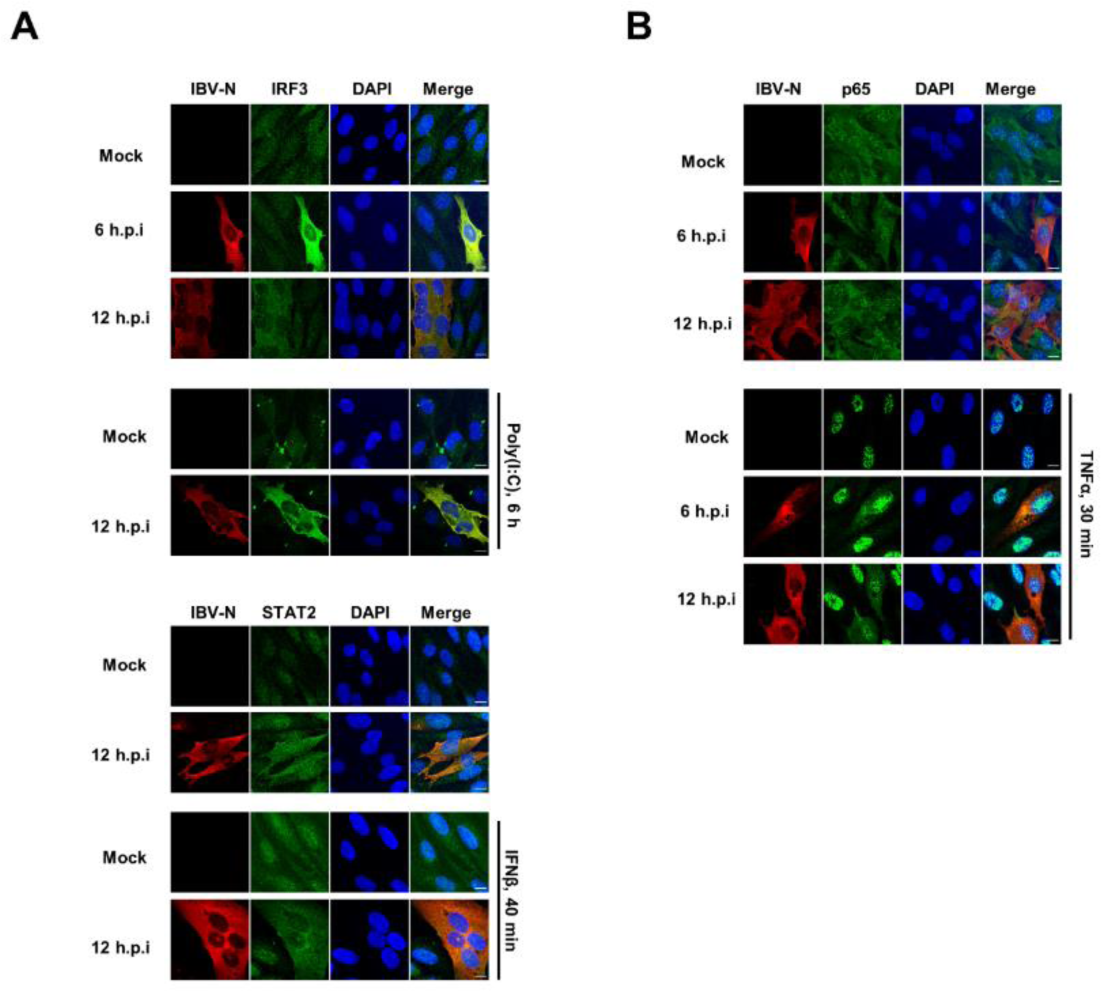
IBV infection suppresses nuclear translocation of transcription factor IRF3, STAT2, and p65 in DF-1 cells. (A-B) DF-1 cells were infected with IBV at an MOI of 1, followed by treatment with poly(I:C), IFNβ, or TNFα. Cells were harvested at the indicated time points and subjected to immunofluorescence analysis. Representative images from three independent experiments are shown. Scale bars: 10 μm.

### IBV Infection induces the phosphorylation of NUP62 and perturbs the integrity of NPC

As IBV infection disrupts the nuclear translocation of multiple transcription factors, we hypothesized that the infection likely interferes with the nucleocytoplasmic trafficking. To investigate this, we analyzed the integrity of NPC in IBV-infected Vero cells by examining the subcellular distribution of the FG-Nups using the monoclonal antibody mAb414. This antibody recognizes a number of FG-Nups, including NUP62, NUP153, NUP98, NUP35, NUP54, NUP58, NUP214, and NUP358 [47], which constitute the mesh in the NPC central channel and are crucial for cargo molecules transport [48]. Immunofluorescence analysis revealed that FG-Nups detected by mAb414 were predominantly localized to nuclear envelope as ring signals; however, in IBV-infected cells, the FG-Nups were dispersed into the cytoplasm at 6 h.p.i. and 12 h.p.i.; they were gradually relocated to the nuclear envelope as ring signals from 12 h.p.i. to 18 h.p.i., coinciding with the formation of large syncytia (Fig 2A, top left panel). These observations suggest that IBV dynamically modulates NPC integrity during infection progression. To corroborate these findings, we performed immunostaining of specific Nups with antibodies against NUP62, NUP153, NUP98, NUP42, and TPR. Results indicated that these specific Nups were primarily localized to the nuclear envelope or inside the nucleus; however, in IBV-infected cells, NUP62, NUP153, NUP42, and TPR were redistributed to the cytoplasm at 6 and 12 h.p.i.; while at 18 h.p.i., NUP42 and TPR were relocated to the nucleus or NE, whereas NUP153 remained dispersed in the cytoplasm and NUP62 was redistributed to one side of the cytoplasm with intense signals (Fig 2A). Compared to other Nups, only a small proportion of NUP98 was dispersed into cytoplasm at 6 h.p.i.; as infected cells formed syncytia at 12 and 18 h.p.i., NUP98 was relocated to nuclear envelope as ring signal. These observations confirm that multiple Nups are dislocated from the nuclear envelope and the NPC integrity is disrupted during early stages of infection (6-12 h.p.i.); at late stage of infection (18 h.p.i), NPC integrity is not fully recovered, as evidenced by the mislocalization of NUP153 and NUP62. Notably, the cytoplasmic dispersion signals of these Nups colocalized well with IBV N protein at 6 and 12 h.p.i., suggesting potential interaction.

Subsequently, we examined the subcellular localization of soluble phase components of nucleocytoplasmic trafficking system: the key receptors importin α1 and importin β1, as well as Ran, which governs the formation and disassembly of transport receptors with cargo proteins. As depicted in Fig 2B, importin β1 exhibited a ring signal adjacent to the NE. However, during IBV infection at 6 h.p.i., a certain proportion of importin β1 was dispersed into the cytosol. Despite this, the majority of this receptor remained associated with the nuclear envelope throughout the infection time course. Importin α1 was predominantly dispersed in the cytoplasm, and IBV infection did not obviously alter its localization. Ran was primarily localized inside the nucleus; however, a small proportion of Ran was translocated to the cytosol and colocalized well with the N protein at 6 and 12 h.p.i. As the infection progressed, Ran relocated to the nucleus at 18 h.p.i.. To validate these results, we further examined the subcellular localization of FG-Nups and Ran in DF-1 cells. Due to the unavailability of antibodies against chicken Nups, only antibodies against human NUP62, NUP98, NUP153, TPR, and Ran, which could cross-react with the corresponding chicken proteins, were applied in this study. Immunofluorescence analysis showed that these proteins were primarily localized to the nucleus or NE; however, in IBV-infected cells, proportions of NUP62, NUP98, NUP153, TPR, and Ran were redistributed from the nuclear envelope or nucleus to the cytoplasm, colocalizing well with N protein (S2A Fig). These results are consistent with the observations in Vero cells. To rule out the possibility of Nups’ cytoplasmic dispersion and their colocalization with the N protein being artifacts resulting from the cross-reaction of two fluorescent secondary antibodies with the same primary antibody, we performed staining for NUP62 and N protein or NUP153 and N protein using a different set of fluorescent secondary antibodies in reverse. The result showed that NUP62 and NUP153 were still dispersed into cytoplasm and colocalized perfectly with IBV N protein (S2B Fig).

**Fig 2.**
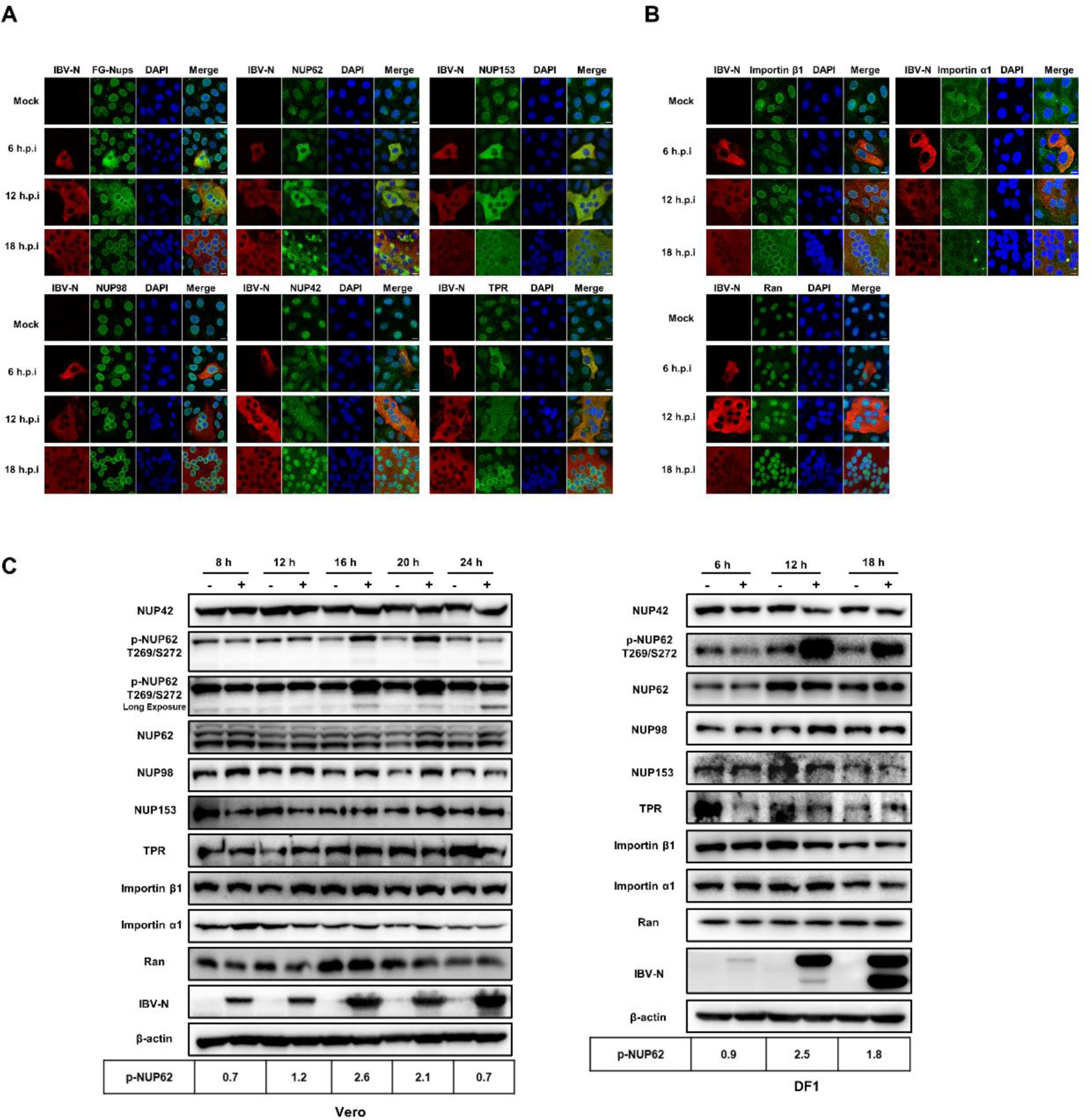
IBV infection disrupts the NPC integrity and induces the phosphorylation of NUP62. (A-B) Vero cells were infected with IBV or mock-infected, harvested at 6, 12, and 18 h.p.i., and subjected to immunofluorescence analysis. Representative images from three independent experiments are shown. Scale bars: 10 μm. (C) Vero and DF-1 cells were infected with IBV or mock-infected, harvested at the indicated time points, and subjected to western blot analysis. β-actin was detected as a loading control. Protein band signals were quantified using Image J, with the intensities of p-NUP62 normalized to total NUP62. The ratio of p-NUP62 in IBV-infected cells to mock-infected cells is presented as p-NUP62 (+:-)

**S2 Fig.**
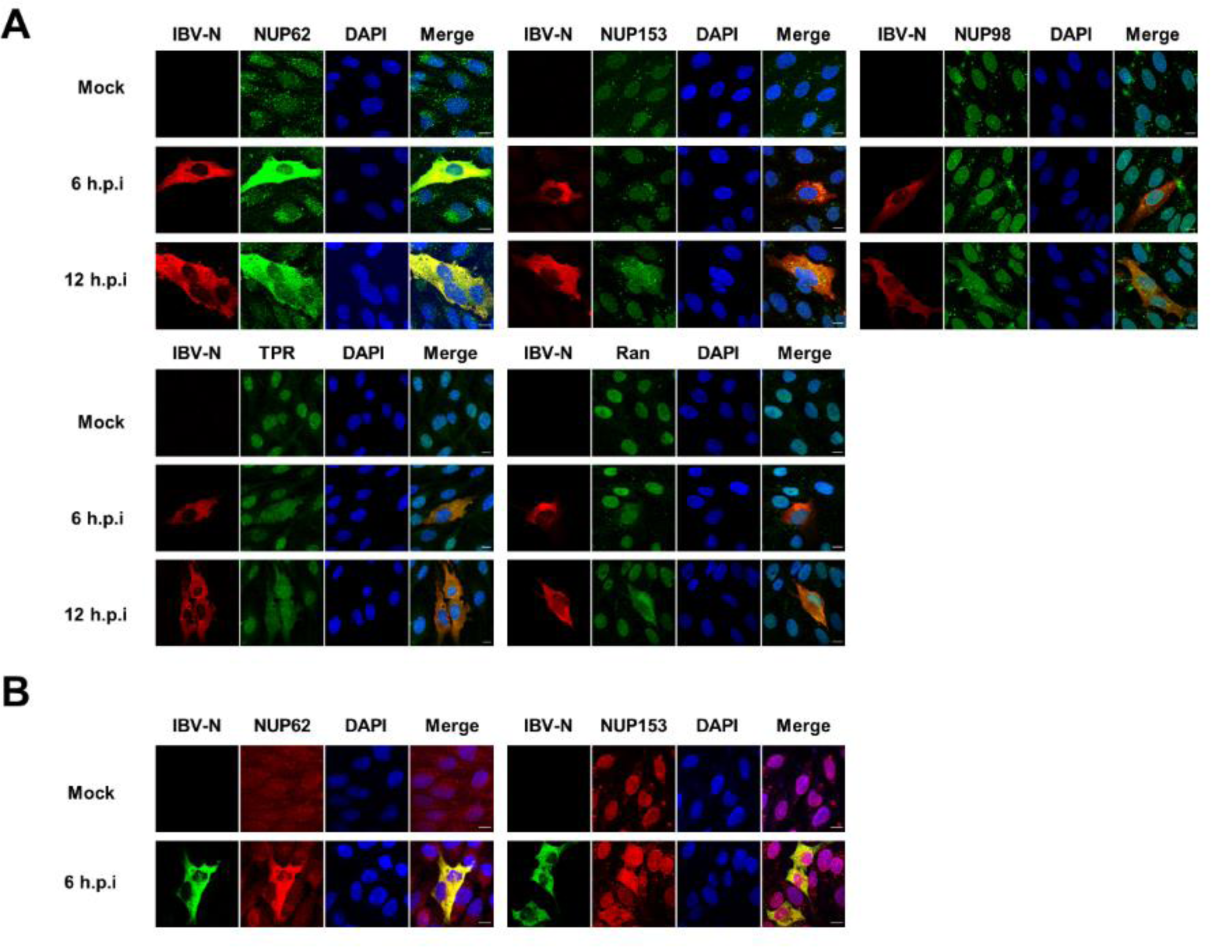
IBV infection induces dislocation of Nups and Ran from nuclear envelope or nucleus to the cytoplasm in DF-1 cells. (A-B) DF-1 cells were infected with IBV at an MOI of 1 or mock-infected, harvested at the indicated time points, and subjected to immunofluorescence analysis. Representative images from three independent experiments are shown. Scale bars: 10 μm.

Next, we examined the expression levels of NUP42, NUP62, NUP98, NUP153, TPR, importin β1, importin α1, Ran, and phosphorylation level of NUP62 (p-NUP62), by western blot analysis. As depicted in Fig 2C, the expression levels of NUP42, NUP62, NUP98, NUP153, importin β1, importin α1, and Ran remained stable in both IBV-infected Vero and DF-1 cells. However, there was a mobility shift observed in NUP42, with the appearance of a smaller band, and phosphorylation of NUP62 at T269 and S272 was observed. Additionally, a potential cleaved p-NUP62 band of approximately 45 kDa was detected in IBV-infected Vero cells. These results suggest that IBV infection alters the post-translational state of NUP42 and NUP62. In summary, these findings suggest that IBV infection primarily disrupts nucleocytoplasmic trafficking function by inducing alterations in the post-translational modifications of Nups and their translocation from the nuclear envelope to the cytoplasm. The redistribution multiple Nups and importin receptors to the cytoplasm likely contributes to the restriction of transcription factor nuclear entry during IBV infection process.

### The IBV N protein is implicated in disrupting nucleocytoplasmic trafficking and hindering the nuclear translocation of transcription factors

To identify viral proteins responsible for the cytoplasmic dispersion of Nups and the obstruction of transcription factor nuclear translocation, IBV proteins were tagged with Flag and expressed in Vero cells. The schematic depiction of IBV-encoded proteins is illustrated in Fig 3A. Immunofluorescence analysis revealed that overexpression of nsp2, nsp5, nsp8, nsp12, nsp13, nsp14, nsp15 and nsp16 did not impact the nuclear envelope localization of FG-Nups; conversely, in cells expressing nsp3, nsp6, nsp7, nsp9, E, M, 5a, and N, mislocalization of FG-Nups was observed (Fig 3A). Subsequently, we investigated whether these viral proteins hindered nuclear translocation of STAT1. As depicted in Fig 3B, nsp2, nsp12, nsp13, E, M and N notably inhibited nuclear translocation of STAT1 induced by IFNβ, nsp5 and nsp6 reduced the STAT1 signal to undetectable levels. Based on these findings, we infer that nsp6, E, M and N not only facilitate the cytoplasmic dispersion of FG-Nups but also impede IFNβ-induced STAT1 nuclear translocation. Notably, among these viral proteins, the N protein prominently promotes the cytoplasmic distribution of FG-Nups, consistent with the observation in IBV-infected cells. Therefore, the IBV N protein emerges as a promising candidate for further investigation. It is noteworthy to mention that the expression of nsp4, nsp10, nsp11, S, 3a, 3b, and 5b was not successful in this study. Therefore, we cannot exclude the possibility that these viral proteins may also contribute to the perturbation of FG-Nups or the inhibition of STAT1 nuclear entry.

**Fig 3.**
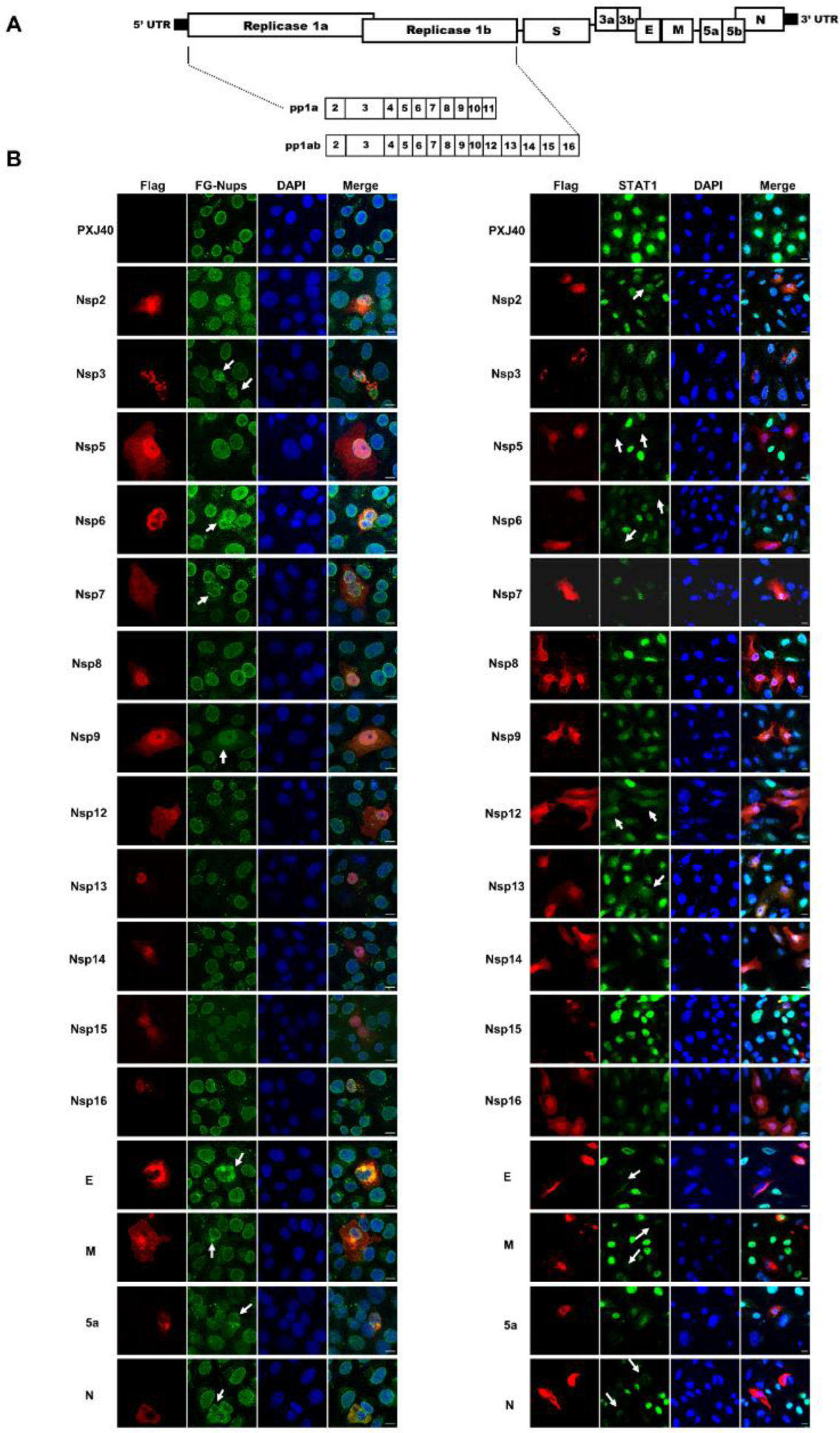
Screening of IBV proteins that induce mislocalization of FG-Nups and inhibit IFNβ-induced nuclear translocation of STAT1. (A) Schematic diagram of the proteins encoded by IBV. (B) Vero cells were transfected with plasmids encoding Flag-tagged IBV proteins or vector PXJ40. At 24 h post-transfection, cells were subjected to immunofluorescence staining for the detection of FG-Nups using mAb414 (left panel). In a parallel group, transfected cells were treated with IFNβ (1000 IU/mL) for 40 min before immunofluorescence staining for detection of STAT1 (right panel). The representative images from three independent experiments are presented. Scale bars: 10 μm.

To further investigate the impact of IBV N protein on the subcellular localization of Nups, Vero cells were transfected with a plasmid encoding IBV N. The subcellular distributions of specific Nups were assessed with corresponding antibodies. Immunofluorescence analysis revealed that in cells expressing the N protein, the subcellular distributions of NUP42, NUP62, and NUP153 were dispersed from the nuclear envelope to the cytoplasm, showing significant colocalization with the N protein. Additionally, the signals of NUP98 and TPR were also altered: NUP98 lost its characteristic intact ring signal and appeared as punctate signals in the cytoplasm, while TPR was dispersed into the cytoplasm with reduced signal intensity (Fig 4A). These findings further support the notion that IBV N is the viral protein responsible for the cytoplasmic dispersion or mislocalization of Nups. Furthermore, importin β1, which is typically localized adjacent to the nuclear envelope as ring signals, exhibited altered morphology associated with the distorted nuclear envelope in cells expressing the N protein (Fig 4B and S3 Fig). Conversely, importin α1, which is primarily dispersed in the cytoplasm, exhibited reduced signals in cells expressing the N protein (Fig 4B and S3 Fig). Regarding Ran, whose signals are predominantly nuclear, they were relocated from the nucleus to the cytoplasm with diminished signal intensity in cells expressing the N protein (Fig 4B). Collectively, these findings provide evidence that the NPC integrity and nucleocytoplasmic trafficking function are perturbed by IBV N protein.

**Fig 4.**
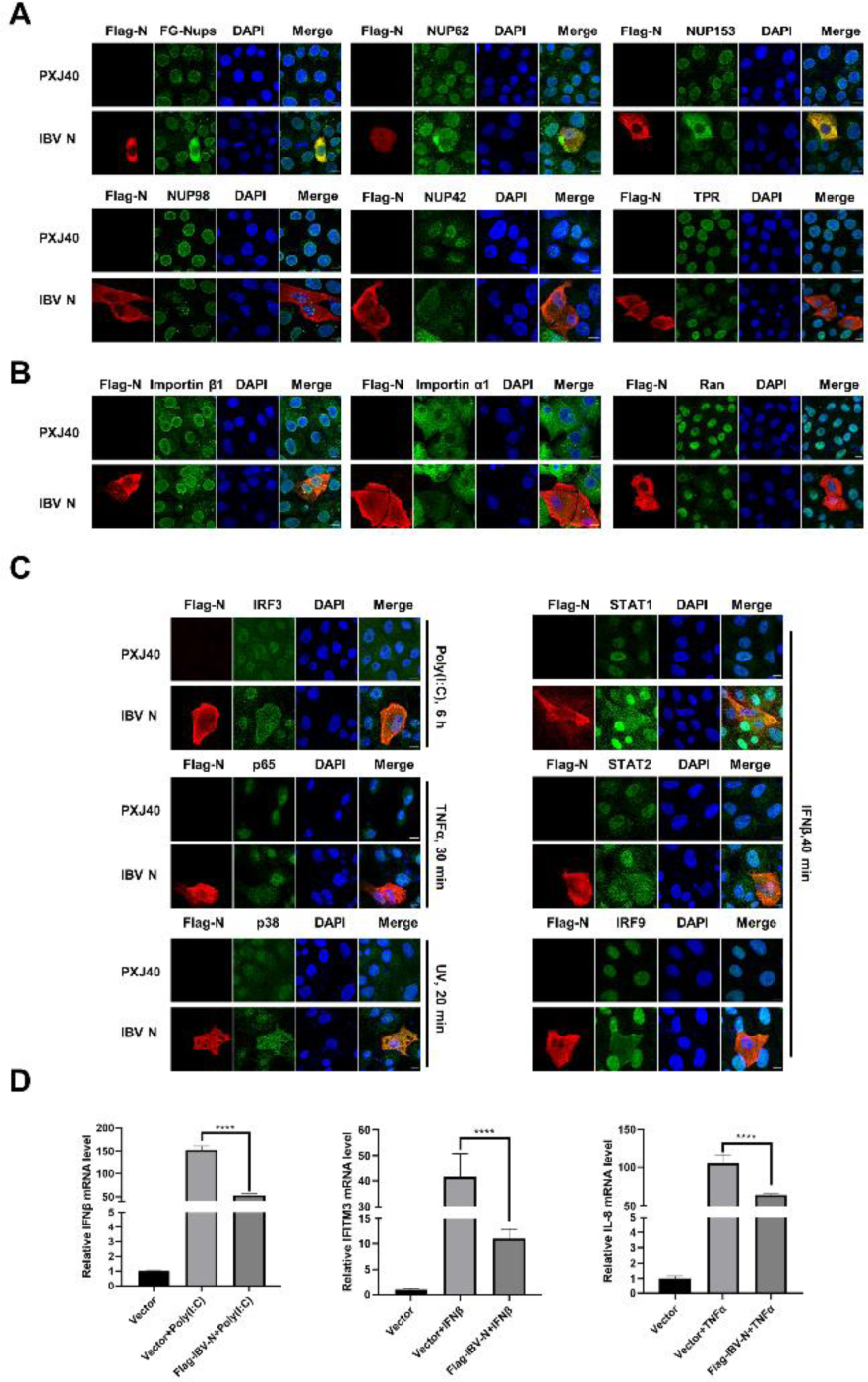
IBV N protein induces dislocation of Nups from the nuclear envelope to the cytoplasm, inhibits nuclear translocation of transcription factors, and suppresses the transcription of antiviral genes. (A-B) Vero cells were transfected with either the vector PXJ40 or a plasmid encoding IBV N protein. At 24 h post-transfection, cells were harvested and subjected to immunofluorescence analysis. (C) Vero cells were transfected with either vector PXJ40 or a plasmid encoding IBV N protein. At 18 h post-transfection, cells were further treated with poly(I:C) (20 μg/mL) for 6 h. In parallel experimental groups, cells were transfected with the vector PXJ40 or a plasmid encoding IBV N protein for 24 h, followed by treatment with IFNβ, TNFα, or UV irradiation, respectively. Cells were then harvested and subjected to immunofluorescence analysis. Representative images from three independent experiments are shown. Scale bars: 10 μm. (D) DF-1 cells were transfected with the vector PXJ40 or a plasmid encoding IBV N protein. At 24 h post-transfection, cells were transfected with poly(I:C) or treated with IFNβ or TNFα for 12 h, followed by qRT-PCR analysis.

**S3 Fig.**
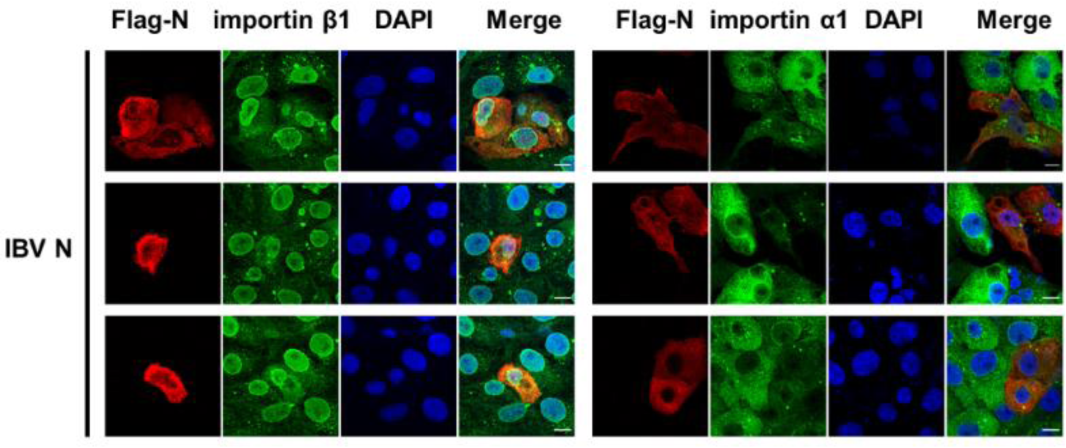
IBV N protein alters the morphology of nuclear envelope and the ring signal of importin β1, and reduces the cytoplasmic signal of importin α1. Vero cells were transfected with either the vector PXJ40 or a plasmid encoding IBV N protein. At 24 h post-transfection, cells were harvested and subjected to immunofluorescence analysis.

To ascertain whether N protein impedes the nuclear translocation of transcription factors, similar to those observations during IBV infection, Vero cells were transfected with the plasmid encoding Flag-tagged IBV N followed by poly(I:C) transfection. According to Fig 4C, the expression of IBV N protein significantly inhibited the poly(I:C)-induced nuclear translocation of IRF3, resulting in the retention of IRF3 in the cytoplasm. Further examination of the JAK-STAT pathway revealed that although IFNβ effectively stimulated the nuclear entry of STAT1, STAT2, and IRF9 in Vero cells, the expression N protein sequestered these transcription factors in the cytoplasm (Fig 4C). Evaluation of NF-κB pathway demonstrated that N protein expression impeded the TNFα-induced nuclear translocation of p65. Similarly, a blockage of UV irradiation-triggered nuclear translocation of p38 MAPK was observed in cells expressing N protein (Fig 4C). Consequently, the transcription of IFNβ, IFITM3, and IL-8, induced by their respective stimuli, was significantly inhibited by the N protein in DF-1 cells (Fig 4D). These results collectively indicate that IBV N protein disrupts NPC integrity and prevents the nuclear translocation of transcription factors, ultimately impairing the expression of downstream antiviral and pro-inflammatory genes.

### The disruption of NPC integrity and interference with nuclear translocation of transcription factors by the N protein is conserved across pan-coronaviruses

The coronavirus N protein is a highly conserved structural protein responsible for encapsulating the viral genomic RNA and facilitating viral RNA synthesis/translation [49]. We aimed to investigate whether the disruption of nucleocytoplasmic trafficking by N protein is conserved across pan-coronaviruses. Flag-tagged N proteins from various genera of coronaviruses, including *alpha-coronavirus* HCoV-H229E, HCoV-NL63, TGEV and PEDV, *beta-coronavirus* MERS-CoV, SARS-CoV, MHV, PHEV, and SARS-CoV-2, and *delta-coronavirus* PDCoV, were expressed in Vero cells, and the subcellular localization of several Nups were examined. As shown in Fig 5A, N proteins from various coronaviruses induced cytoplasmic dispersion of FG-Nups, NUP62, NUP153, and NUP42, with significant colocalization with the N protein. TPR exhibited a cytoplasmic dispersion pattern with reduced signal intensity in all N protein-expressing cells. Conversely, NUP98 lost its intact ring signal and formed punctate aggregates in the cytoplasm in the majority of N protein-expressing cells (including those from HCoV-NL63, MERS-CoV, SARS-CoV, MHV, PHEV, SARS-CoV-2, and PDCoV). The subcellular localization of two importin receptors and Ran was also assessed. As illustrated in Fig 5B, in all N protein-expressing cells, importin β1 lost its intact ring signals on the nuclear envelope and displayed as mislocated signal. Additionally, a decreased signal of importin α1 was observed, and Ran was dispersed into the cytoplasm with diminished signal. Taken together, these data suggest that the ability to dismantle the NPC and interfere with nucleocytoplasmic trafficking is conserved across pan-coronaviruses N proteins.

**Fig 5.**
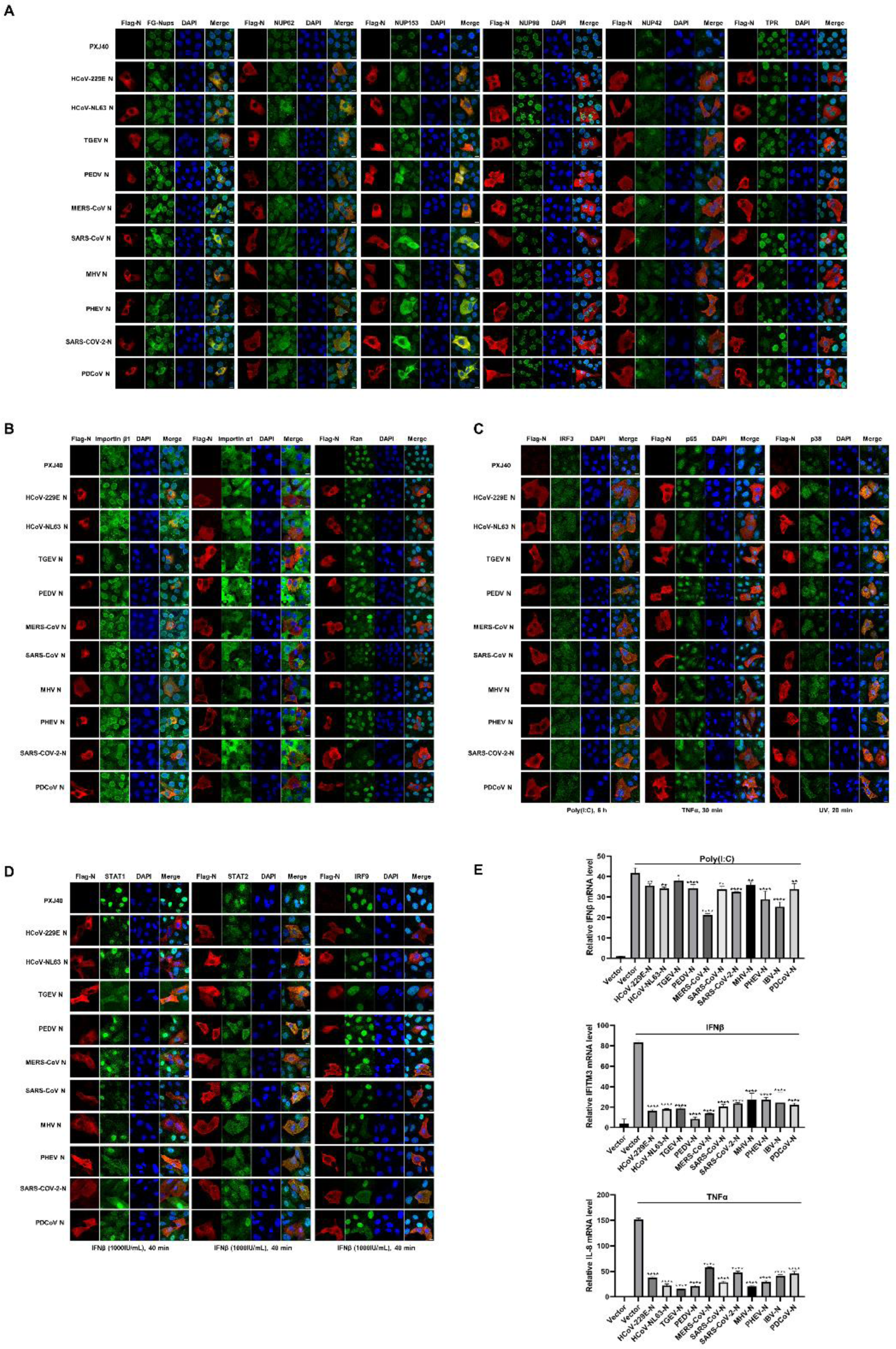
Disruption of nucleocytoplasmic trafficking, inhibition of nuclear translocation of transcription factors, and suppression of antiviral genes transcription by pan-coronaviruses N proteins. (A-B) Vero cells were transfected with plasmids encoding Flag-tagged N proteins from different genera of coronaviruses, while vector PXJ40 served as a control. Cells were harvested 24 h post-transfection and subjected to immunofluorescence analysis. (C-D) Vero cells were transfected with plasmids encoding N proteins or vector PXJ40. At 18 h post-transfection, cells were treated with poly(I:C), IFNβ, TNFα, or UV irradiation, respectively, followed by immunofluorescence analysis. Representative images from three independent experiments are shown. Scale bars: 10μm. (E) HEK-293T cells were transfected with plasmids encoding N proteins or vector PXJ40. At 24 h post-transfection, cells were treated with poly(I:C), IFNβ, or TNFα. Cells were harvested at 12 h post-treatment, and the transcription levels of IFNβ, IFITM3, or IL-8 were detected using qRT-PCR.

Subsequently, we investigated whether the nuclear translocation of transcription factors is impeded by N proteins from various coronaviruses. Vero cells were transfected with N proteins, followed by poly (I:C) transfection. As shown in Fig 5C-D, poly(I:C) successfully induced the nuclear translocation of IRF3; however, in all N protein-expressing cells, IRF3 remained in the cytoplasm. Further examination of the JAK-STAT pathway also revealed that, although IFNβ successfully stimulated the nuclear translocation of STAT1, STAT2, and IRF9, in all N protein-expressing cells, STAT1, STAT2, and IRF9 were dispersed in the cytoplasm. Moreover, the transcription factors p65 and p38 MAPK entered the nucleus following TNFα treatment or UV irradiation; however, the expression of N proteins impeded the nuclear translocation of these transcription factors. Alongside these observations, the transcription of IFNβ, IFITM3 and IL-8 in response to corresponding stimuli were significantly inhibited by all N proteins from different coronaviruses (Fig 5E). Overall, these findings demonstrate the blockage of multiple transcription factors’ nuclear translocation and inhibition of antiviral gene expression by the N protein is conserved across pan-coronaviruses.

### Pan-coronaviruses N proteins interact with scaffold protein RACK1

To delve into the mechanisms underlying how the N protein modulates the nucleocytoplasmic trafficking, we conducted co-immunoprecipitation (Co-IP) assays combined with liquid chromatography and mass spectrometry (LC-MS/MS) to screen cellular proteins interacting with IBV N protein. A comprehensive analysis identified a total of 694 cellular candidates that co-immunoprecipitated with IBV N protein. Gene ontology (GO) annotation and Kyoto Encyclopedia of Genes and Genomes (KEGG) database analyses revealed a significant enrichment of host gene expression pathways. These include mitochondrial translation, mRNA metabolic process, ribosome biogenesis, regulation of translation, ribonucleoprotein complex assembly, RNA localization, spliceosomal complex assembly, RNA modification, RNA 3’-end processing, among others (Fig 6A-B). Notably, these findings align with the previous interactome analyses of the N proteins from IBV [50], PEDV [51], SARS-CoV-2 [52, 53]. Among the IBV N binding proteins involved in nucleocytoplasmic transport, key players were identified and listed in Fig 6C, including RACK1 (Receptor of activated protein C kinase 1), NXF1 (Nuclear RNA export factor 1), LMNB1 (Lamin B1), KPNA2 (importin α1), PPP1CC (Protein phosphatase PP1γ), RAE1 (mRNA export factor), LBR (Lamin B receptor), SRPK1 (SRSF protein kinase 1), NUPL2 (Nucleoporin-like protein 2, NUP42).

**Fig 6.**
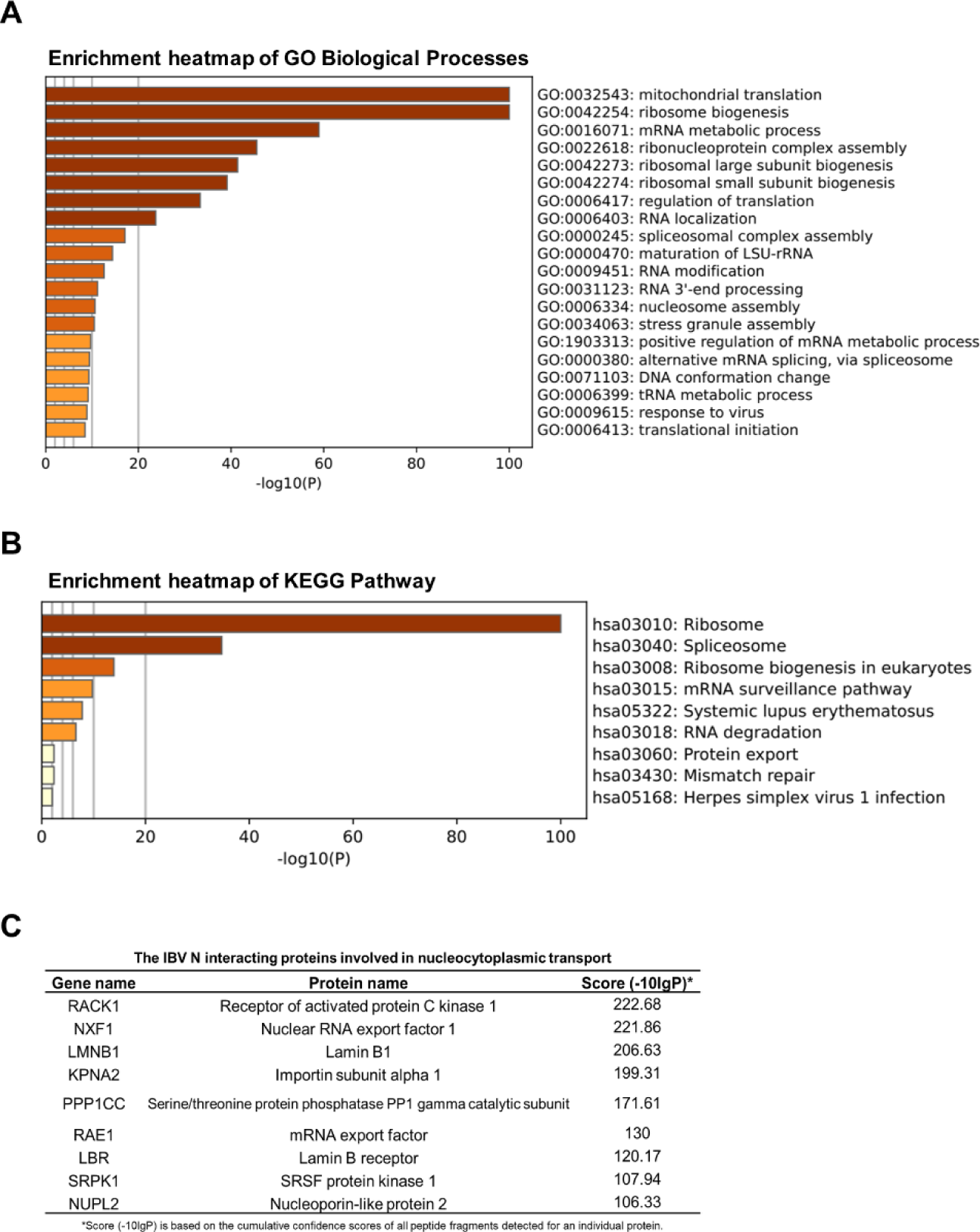
Bioinformatic analysis of IBV N-protein interactome. Plasmid encoding IBV N protein was transfected into HEK-293T cells for 24 h. Whole-cell lysates were immunoprecipitated using anti-Flag antibody, and three independent immunoprecipitated samples were subjected to LC-MS/MS analysis. PXJ40-transfected cell lysates served as negative control to eliminate nonspecific binding proteins. (A) Heatmap illustrating the enrichment of Gene Ontology (GO) annotations within the IBV N-protein interactome. (B) Heatmap displaying the enrichment of Kyoto Encyclopedia of Genes and Genomes (KEGG) pathways in the IBV N-protein interactome. (C) IBV N protein interacts with several cellular proteins involved in nucleocytoplasmic transport.

Among the proteins interacting with IBV N, as listed in Fig 6C, RACK1 stands out as highly conserved intracellular adaptor protein with pivotal roles in anchoring and stabilizing proteins activity, and shuttling proteins to specific cellular location, such as activated PKC [54]. To validate the interaction between N protein and RACK1, we co-transfected plasmids encoding Flag-tagged IBV N and HA-tagged RACK1 into HEK-293T cells. The specific binding between IBV N protein and RACK1 was confirmed by Co-IP using anti-HA or anti-Flag antibodies: both antibodies successfully precipitated Flag-N and HA-RACK1 together (Fig 7A). We further examined whether IBV N binds to endogenous RACK1 during virus infection. DF-1 cells were infected with IBV, followed with immunoprecipitation using anti-IBV N polyclonal antibody. The results demonstrated that the anti-IBV N antibody efficiently co-precipitated N protein and endogenous RACK1, providing evidence of specific binding during the virus infection process (Fig 7B). Since the antibody against human RACK1 recognizes the linear epitopes of chicken RACK1 but not its conformational epitopes, DF-1 cells are not suitable for studying the subcellular localization of endogenous RACK1. Therefore, Vero cells were employed for subsequent immunofluorescence study. The results in Fig 7C revealed that in mock-infected Vero cells, endogenous RACK1 was distributed in both the nucleus and cytoplasm. However, in IBV-infected Vero cells, a proportion of endogenous RACK1 was dispersed into the cytoplasm at 6 and 12 h.p.i., where it co-localized with IBV N protein, indicating alterations in its localization by N protein during infection.

**Fig 7.**
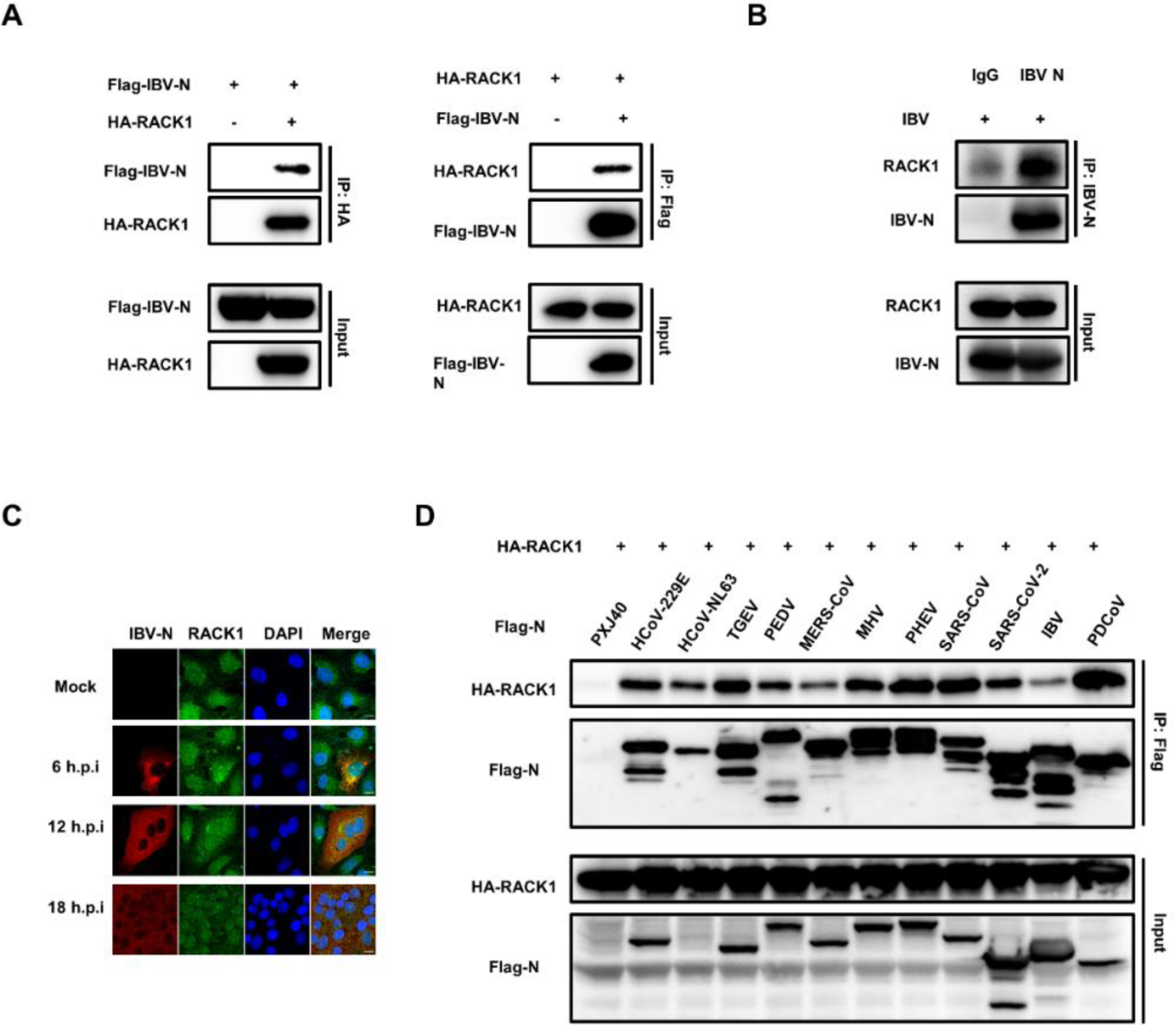
Pan-coronaviruses N proteins interact with RACK1. (A) HEK-293T cells were co-transfected with plasmids encoding Flag-IBV N and HA-RACK1. Co-transfections with Flag-IBV-N and PCMV-HA, or PXJ40 and HA-RACK1 were performed as control groups. Cell lysates were immunoprecipitated using anti-Flag or anti-HA antibodies, followed by immunoblot analysis. (B) DF-1 cells were infected with IBV at an MOI of 1. At 12 h.p.i., cell lysates were immunoprecipitated using anti-IBV N antibody, followed by immunoblot analysis. (C) Vero cells were infected with IBV at an MOI of 1 or mock-infected. Immunostaining was performed at 6, 12, and 18 h.p.i. Representative images from three independent experiments are shown. Scale bars: 10 μm. (D) HEK-293T cells were co-transfected with plasmids encoding Flag-tagged N proteins from different coronaviruses and HA-RACK1. Co-transfection of PXJ40 with plasmid encoding HA-RACK1 served as a control. Cell lysates were immunoprecipitated using anti-Flag antibody, followed by immunoblot analysis.

To investigate whether the interaction with RACK1 is a common feature among pan-coronaviruses N proteins, Flag-tagged N proteins from various coronaviruses were co-expressed with HA-RACK1 in HEK-293T cells. Co-IP results showed that the anti-Flag antibody efficiently precipitated both Flag-N proteins and HA-RACK1, while no RACK1 was pulled down in the absence of N protein (PXJ40 transfection group) (Fig 7D), indicating that the interaction with RACK1 is a conserved characteristic among pan-coronaviruses N proteins.

### The cytoplasmic redistribution of Nups requires PKCα/β activity

RACK1 serves as an anchoring protein for PKC and is responsible for trafficking PKC to specific subcellular locations. Although PKC has been reported to be involved in the modulation of lamin B1 phosphorylation and the cell cycle [55–57], whether PKC phosphorylates Nups remains unexplored. Notably, phosphorylation and cytoplasmic distribution of NUP62 were observed in IBV-infected cells, and N protein was found to be responsible for the cytoplasmic distribution of Nups. Intriguingly, N protein interacts with the PKC scaffold protein RACK1. These observations prompt us to speculate that N protein might regulate PKC activity via RACK1 and be involved in NUP62 phosphorylation. To test our hypothesis, we first examined whether PKC activity is implicated in the cytoplasmic distribution of Nups induced by N protein expression. Vero cells were transfected with Flag-tagged N proteins from different coronaviruses and the subcellular localization of PKCα/β, FG-Nups, NUP62, and NUP153 was examined. Immunofluorescence analysis revealed that PKCα/β translocated from the nucleus to the cytoplasm together with FG-Nups, NUP62, or NUP153 in all N protein-expressing cells (Fig 8A), demonstrating that N protein alters the subcellular localization of PKCα/β and affects their activity. The pronounced colocalization of N, PKCα/β, and FG-Nups/NUP62/NUP153 suggests potential interactions among N kinases and Nups. The application of the PKCα/β inhibitor Enzastaurin resulted in the majority of PKCα/β, FG-Nups, NUP62, and NUP153 relocating to the nucleus or perinuclear region, while all N proteins remained diffusely distributed throughout the cytoplasm (Fig 8A). Thus, the activity of PKCα/β is implicated in the cytoplasmic dispersion of Nups in N protein-expressing cells.

**Fig 8.**
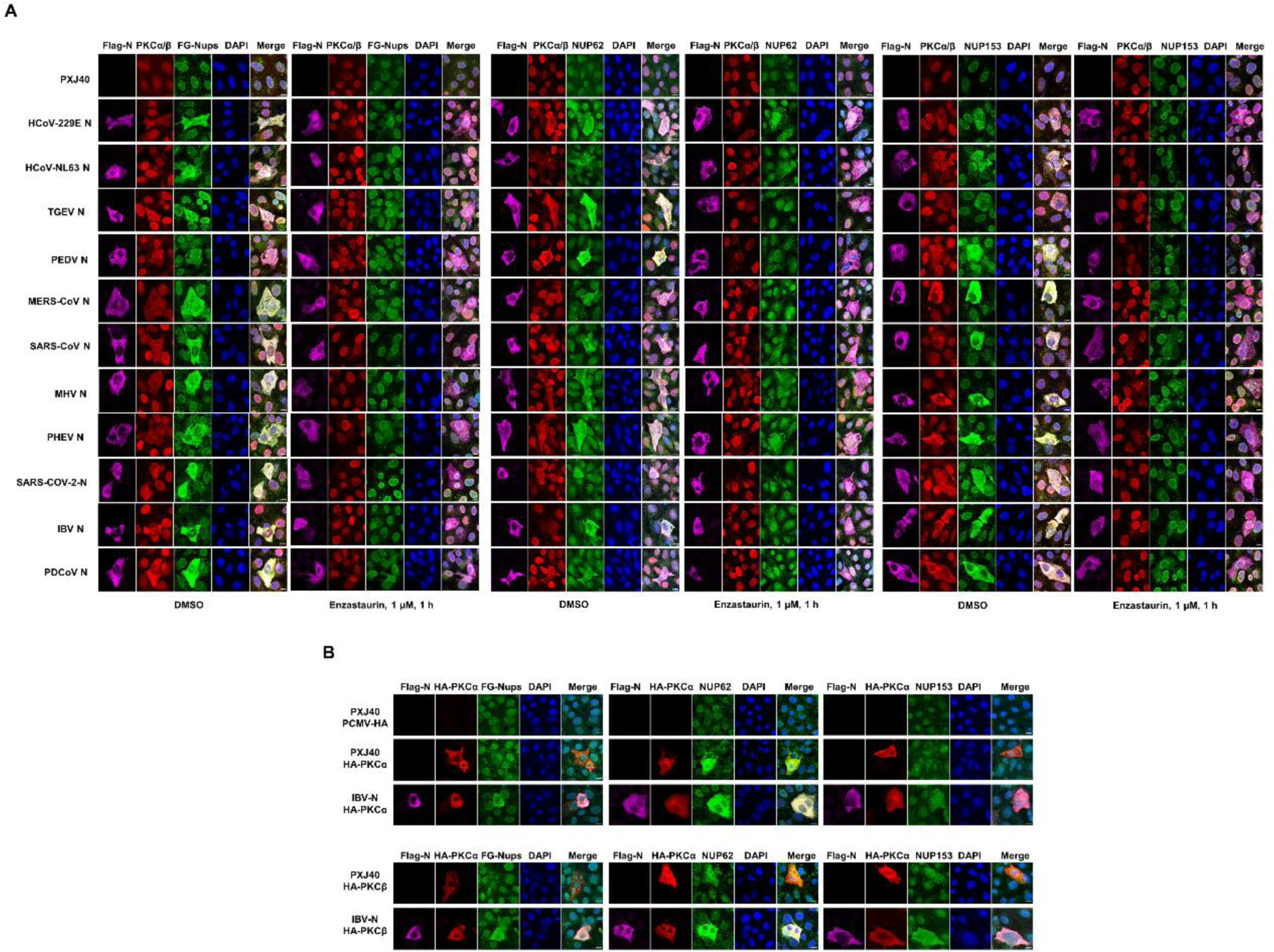
Implication of PKCα/β activity in the cytoplasmic distribution of Nups. (A) Vero cells were transfected with plasmids encoding Flag-tagged N proteins from different coronaviruses or PXJ40. After 24 h post-transfection, cells were treated with Enzastaurin (1 μM) or DMSO for 1 h, followed by immunostaining to assess the subcellular distribution of N, PKCα/β, and FG-Nups/NUP62/NUP153. (B) Plasmid encoding HA-tagged PKCα or PKCβ was co-transfected with PXJ40, or co-transfected with plasmid encoding Flag-tagged IBV N for 24 h, followed by immunostaining to evaluate the subcellular localization distribution of PKCα, PKCβ, N, FG-Nups, NUP62, and NUP153. Representative images from three independent experiments are presented. Scale bars: 10 μm.

To further confirm the implication of PKC in the cytoplasmic distribution of Nups, HA-tagged PKCα and PKCβ were expressed in Vero cells alone or together with IBV N protein. Immunofluorescence analysis revealed that overexpression of both HA-PKCα and HA-PKCβ promoted the cytoplasmic distribution of NUP62, while the cytoplasmic dispersion of FG-Nups and NUP153 was less pronounced. Surprisingly, co-expression of IBV N protein and PKCα/PKCβ led to enhanced cytoplasmic distribution of FG-Nups, NUP62, and NUP153, with these Nups colocalizing well with N and PKCα/PKCβ (Fig 8B). This observation suggests that PKCα and PKCβ are capable of inducing the cytoplasmic dispersion of NUP62, and the presence of N protein promotes the dispersion of more Nups into the cytoplasm.

### The activity of PKCα/β is implicated in the phosphorylation of NUP62 and the cytoplasmic distribution of Nups during IBV infection

To determine whether IBV infection activates PKC activity, we evaluated the phosphorylation levels of PKCα and PKCβ in IBV-infected Vero cells and DF-1 cells. As illustrated in Fig 9A, compared to mock-infected cells, the phosphorylation level of PKCβ (p-PKCβ, at S660) gradually increased over the course of infection, peaking at 18 h.p.i., while the phosphorylation level of PKCα (p-PKCα, at T638) remained stable. Remarkably, the level of p-NUP62 (at T269/S272) also exhibited a gradual increase during IBV infection, concurrent with PKCβ phosphorylation. Immunofluorescence analysis in Fig 9B revealed that during the early stages of infection (6 and 12 h.p.i.), PKCα/β translocated from the nucleus to the cytoplasm and exhibited diffuse colocalization with N protein. However, as the infection progressed, PKCα/β gradually relocated to the nucleus (12-18 h.p.i.), indicating dynamic regulation of their subcellular localization by IBV infection. The alteration in subcellular localization of PKCα/β induced by virus infection likely determines their proximity to substrates, influencing their kinase activity.

**Fig 9.**
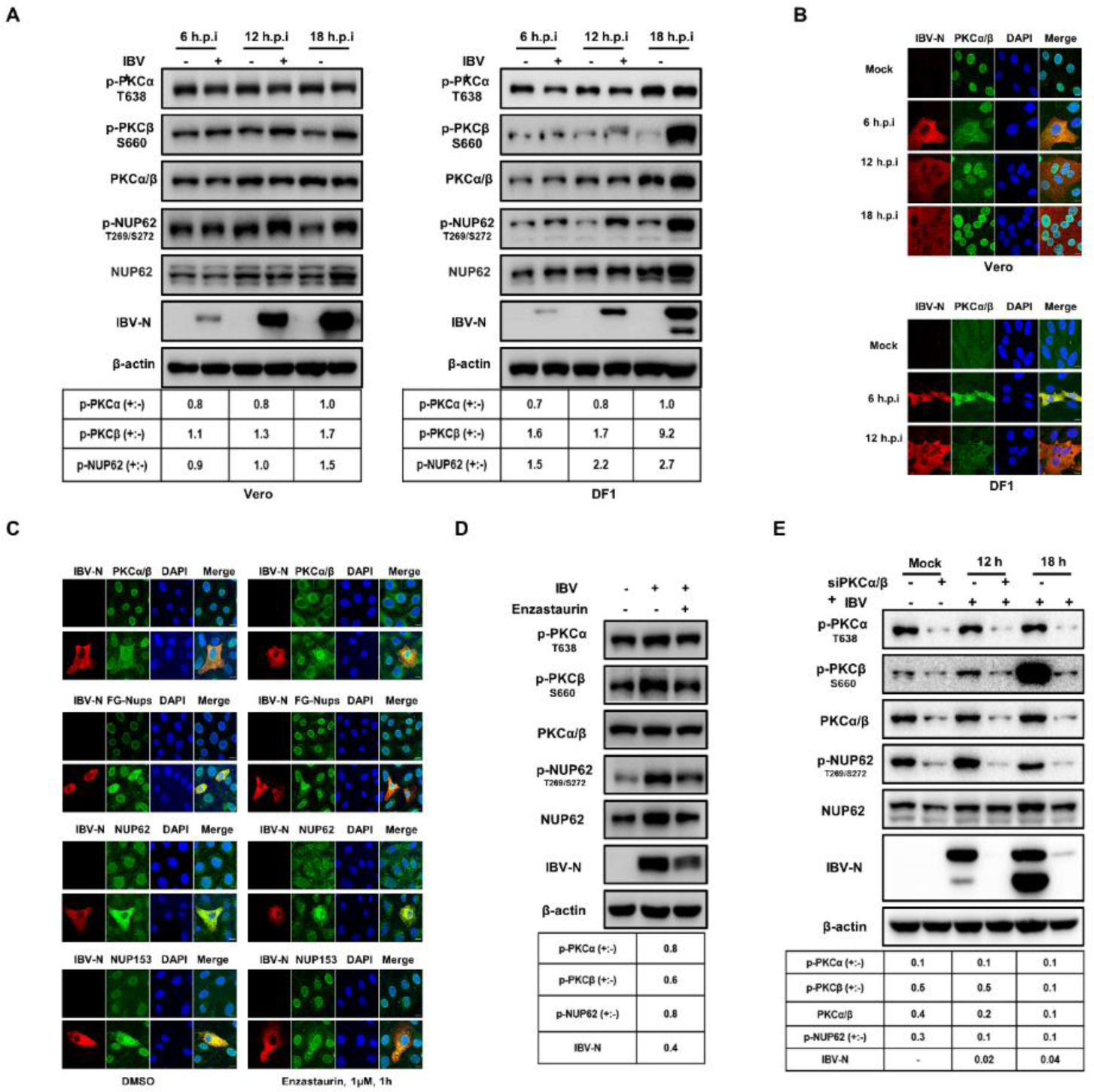
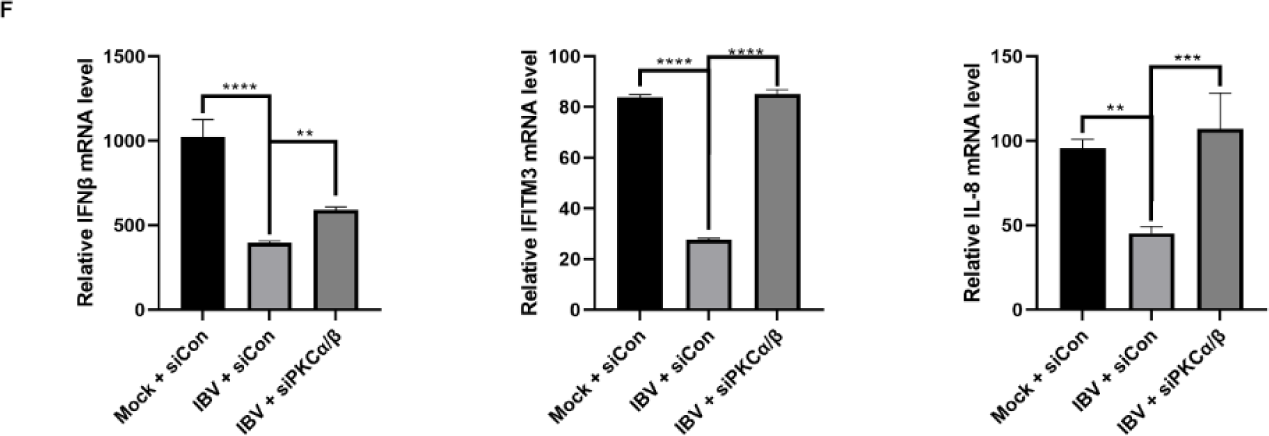
role of PKCα/β activity in NUP62 phosphorylation, the cytoplasmic distribution of FG-Nups, suppression of the antiviral response, and virus replication. (A) Vero or DF-1 cells were infected with IBV at an MOI of 1 or mock-infected. Cells were harvested at 6, 12, and 18 h.p.i. and subjected to western blot analysis using corresponding antibodies. The intensities of p-PKCα, p-PKCβ, and p-NUP62 bands were normalized to total PKCα/β or NUP62, respectively. The ratio of p-PKCα, p-PKCβ, and p-NUP62 in IBV-infected cells to those in mock-infected cells is denoted as p-PKCα (+:-), p-PKCβ (+:-), and p-NUP62 (+:-). (B) Vero and DF-1 cells were infected with IBV or mock-infected and subjected to immunostaining. (C) Vero cells were infected with IBV or mock-infected. At 2 h.p.i., cells were treated with DMSO or Enzastaurin (1μM) for 4 h. The subcellular localization of the indicated proteins was analyzed with immunostaining. Representative images from three independent experiments are presented. Scale bars: 10 μm. (D) DF-1 cells were infected with IBV or mock-infected, followed by treatment with DMSO or Enzastaurin (1μM) at 18 h.p.i. Western blot analysis was performed with indicated antibodies. (E) DF-1 cells were transfected with siRNA targeting PKCα/β for 48 h, followed by IBV infection. Western blot analysis was conducted to detect the indicated proteins. The ratio of p-PKCα, p-PKCβ, p-NUP62, and N protein in Enzastaurin-treated cells or siPKCα/β-transfected cells to those in DMSO-treated or siControl-transfected cells is denoted as p-PKCα (+:-), p-PKCβ (+:-), NUP62 (+:-), and N (+:-). (F) DF-1 cells were transfected with siRNA targeting PKCα/β for 48 h, followed by IBV infection at an MOI of 5. The infected cells were treated with poly(I:C) or TNFα at 2 h.p.i for 6 h, or subjected to IFNβ treatment at 6 h.p.i for 6 h. qRT-PCR was performed to detect the transcription levels of IFNβ, IFITM3, and IL-8.

The next question we addressed was whether PKCα/β activity is implicated in the cytoplasmic dispersion of Nups during IBV infection. We treated IBV-infected cells with the PKCα/β inhibitor Enzastaurin and examined the subcellular localization of Nups. As shown in Fig 9C, compared to the DMSO-treated group, Enzastaurin treatment caused PKCα/β, FG-Nups, NUP62, and NUP153 to relocate to the nucleus and perinuclear region in IBV-infected cells, resulting in reduced cytoplasmic dispersion. These observations suggest that PKCα/β activity is necessary for IBV infection-induced Nups cytoplasmic redistribution, similar to what was observed in N protein-expressing cells. Interestingly, Enzastaurin treatment reduced the phosphorylation levels of PKCβ, NUP62, and the expression of IBV N protein (Fig. 9D), indicating that the activation of PKCα/β is involved in NUP62 phosphorylation and benefits virus replication.

The impact of PKCα/β on NUP62 phosphorylation was further investigated by suppressing PKCα and PKCβ expression using a siRNA targeting both PKCα and PKCβ. As depicted in Fig 9E, transfection with PKCα/β siRNA effectively suppressed the expression of both PKCα and PKCβ, leading to a substantial reduction in total PKCα/β levels, as well as p-PKCα and p-PKCβ. Concurrently, phosphorylation of NUP62 (p-NUP62) decreased to nearly undetectable levels in both mock- and IBV-infected cells, indicating the essential role of PKCα/β in NUP62 phosphorylation, under both normal physiological conditions and during virus infection. Furthermore, depletion of PKCα/β resulted in a significant decrease in IBV N protein levels, underscoring the necessity of PKCα/β for efficient virus replication.

To investigate whether phosphorylation of NUP62 is responsible for its cytoplasmic distribution, we treated Vero cells with the PP1 and PP2A inhibitor okadaic acid to inhibit dephosphorylation events. We examined the phosphorylation levels of PKCα, PKCβ, and NUP62 by Western blot analysis. As shown in S4A Fig, okadaic acid treatment led to a slight accumulation of p-PKCα, accompanied by the appearance of two additional bands corresponding to 110 or 180 kDa, which represent dimers/trimers with enhanced activity [58, 59]. The phosphorylation form of PKCβ (p-PKCβ) was also greatly accumulated. Therefore, PP1 or PP2A is implicated in the dephosphorylation of PKCα/PKCβ and suppression of their activity. Simultaneously, the phosphorylation form of NUP62 (p-NUP62) was predominantly accumulated. Meanwhile, a band with slower mobility at approximately 120 kDa was detected, which might represent a potential dimer. The dimerization or oligomerization of p-NUP62 is involved in self-interaction or interaction with other Nups [60, 61], although the underlying mechanism and functional consequence are unclear. Total NUP62 exhibited minor mobility shift bands representing hyperphosphorylation forms at multiple sites. The significant accumulation of p-NUP62 by okadaic acid treatment demonstrates that PP1 or PP2A directly dephosphorylates NUP62 and probably contribute to the disassembly of NPC. Immunofluorescence analysis showed that okadaic acid treatment greatly promoted the cytoplasmic distribution of NUP62 (S4B Fig, top panel), confirming the correlation between phosphorylation of NUP62 and its cytoplasmic dispersion. Additionally, a small proportion of FG-Nups was also dispersed to the cytosol in okadaic acid-treated cells (S4B Fig, lower panel), further supporting the idea that phosphorylation events lead to the dissociation of FG-Nups.

**S4 Fig.**
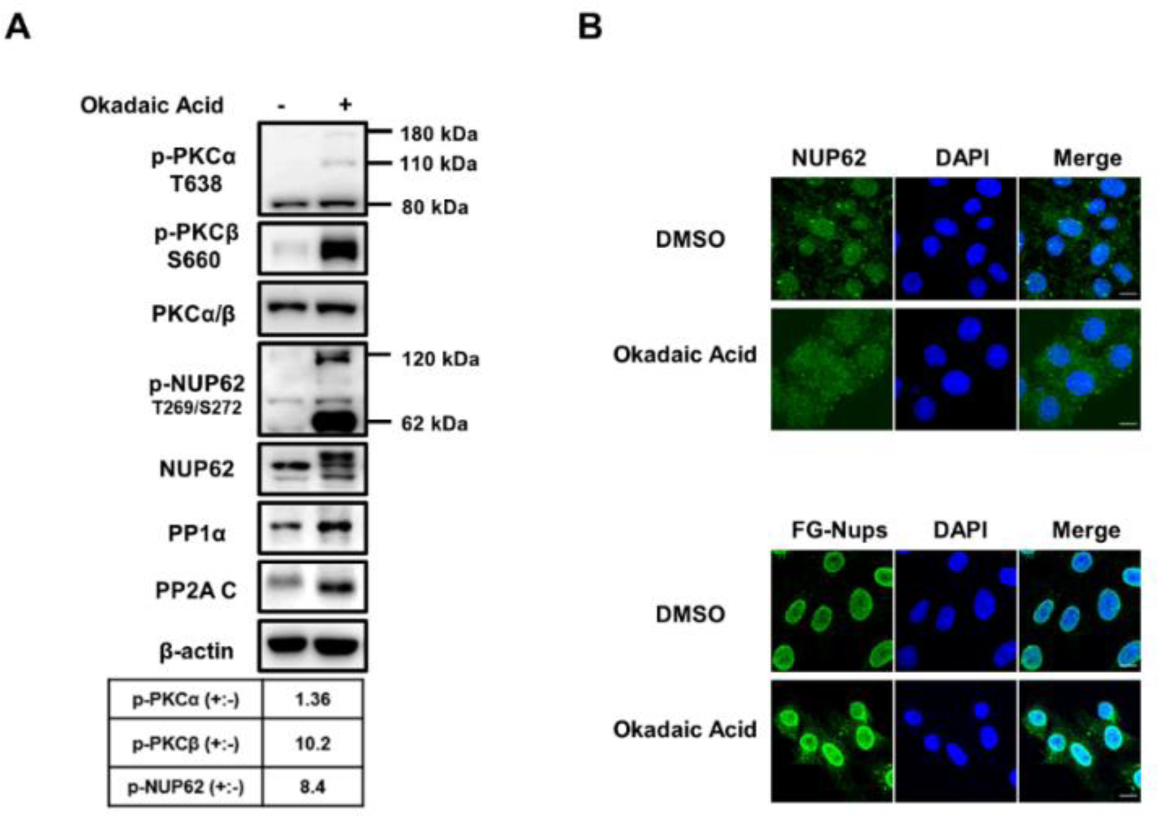
Effect of PP1 and PP2A inhibitor okadaic acid treatment on phosphorylation of PKCα, PKCβ and NUP62, and induction of cytoplasmic dispersion of NUP62 and FG-Nups. (A) DF-1 cells were treated with either DMSO or okadaic acid (1 μM) for 1 h and subjected to western blot analysis using the indicated antibodies. The intensities of p-PKCα, p-PKCβ, and p-NUP62 bands were normalized to total PKCα/β or NUP62. The ratio of p-PKCα, p-PKCβ, and p-NUP62 in okadaic acid-treated cells to DMSO-treated cells is denoted as p-PKCα (+:-), p-PKCβ (+:-), and p-NUP62 (+:-). (B) Vero cells were treated with either DMSO or okadaic acid (1 μM) for 1 h and subjected to immunostaining. Representative images are shown. Scale bars: 10 μm.

Finally, we examined whether depletion of PKCα/β affects the expression profile of antiviral genes. Real-time qRT-PCR results revealed that IBV infection compromised the expression of IFNβ, IFITM3, and IL-8 induced by corresponding stimuli. In PKCα and PKCβ knockdown cells, the transcription of these antiviral genes was significantly restored (Fig 9F). This observation suggests that PKCα/β is involved in the suppression of the innate immune response during IBV infection.

Overall, the above results demonstrate that IBV infection activates PKCβ through phosphorylation and synchronously alters the subcellular localization of PKCα/β. Meanwhile, the activated PKCα/β is responsible for phosphorylating NUP62 and distributing Nups to the cytoplasm, ultimately suppressing the expression of antiviral genes and facilitating virus replication. However, due to the indiscriminate knockdown or inhibition of PKCα and PKCβ by siRNA or chemical inhibitors, we are unable to determine from the current data whether PKCα or PKCβ specifically mediates the phosphorylation of NUP62.

### RACK1 is essential for the phosphorylation of PKCα/β and NUP62, the suppression of antiviral gene expression, and the promotion of IBV infection

To investigate whether RACK1 is involved in the regulation of PKC activity and NUP62 phosphorylation, we knocked down RACK1 in DF-1 cells, followed by IBV infection. As illustrated in Fig 10A, the reduced expression of RACK1 led to a decrease in p-PKCα, p-PKCβ, and p-NUP62 levels in both mock- and IBV-infected DF-1 cells, demonstrating that RACK1 participates in the phosphorylation of PKCα, PKCβ, and NUP62. Concurrently, the expression level of IBV N protein was also reduced in RACK1 knockdown cells, revealing the importance of RACK1 in virus infection. Furthermore, depletion of RACK1 significantly restored the transcription of IFNβ, IFITM3, and IL-8 in IBV-infected DF-1 cells, under conditions with or without chemical stimuli (Fig 10B-C). These results demonstrate that RACK1 is an essential host factor for PKCα and PKCβ to maintain and exert kinase activity on phosphorylating NUP62, thereby suppressing the expression of antiviral genes and ultimately benefiting IBV replication.

**Fig 10.**
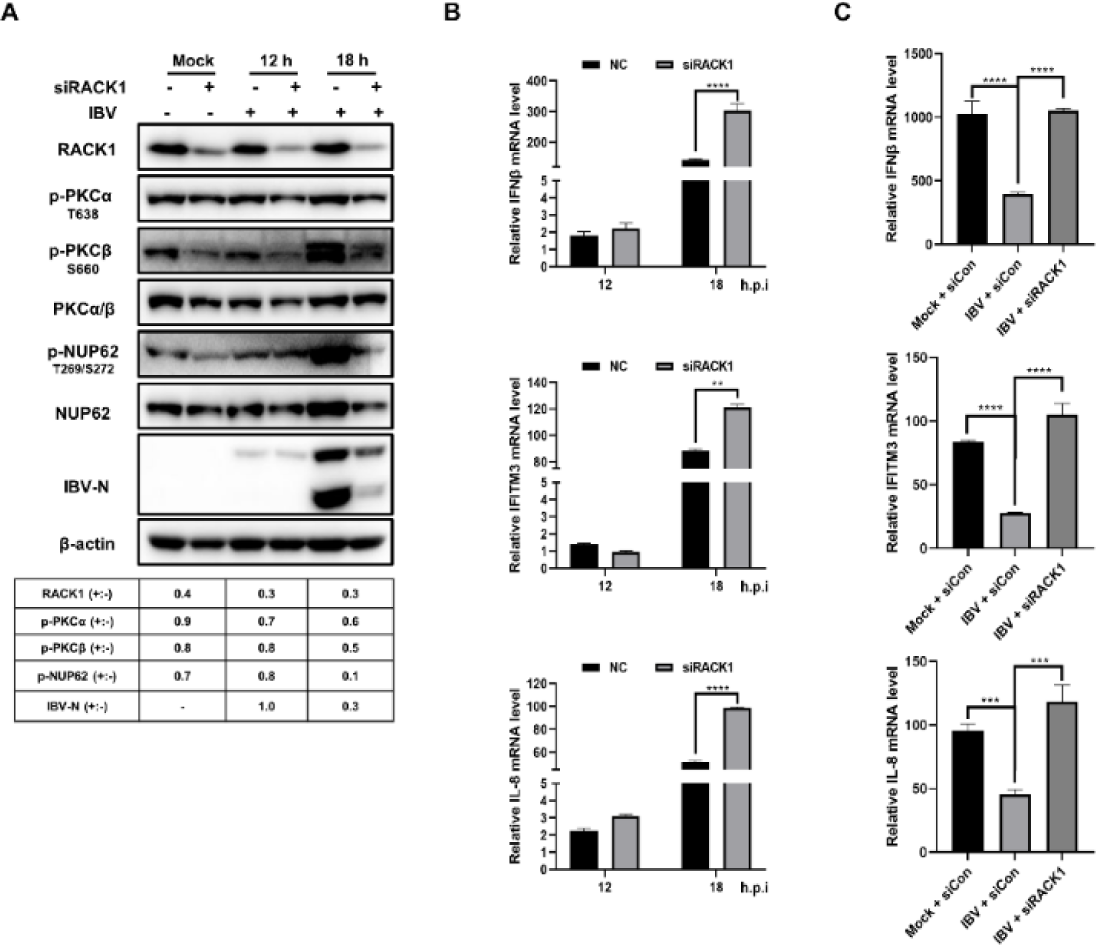
Role of RACK1 in maintaining phosphorylation of PKCα, PKCβ, and NUP62, suppression of antiviral gene expression, and promotion of IBV infection. (A-C) DF-1 cells were transfected with RACK1 siRNA or control siRNA, followed by IBV infection at 48 h post-transfection. One parallel experimental group was subjected to treatment with poly(I:C) transfection, TNFα, or IFNβ, respectively. Cells were harvested at indicated time points and subjected to Western blot analysis (A) or qRT-PCR analysis (B and C). For panel A, the intensities of RACK1, p-PKCα, p-PKCβ, p-NUP62, and IBV N bands were normalized to β-actin, PKCα/β, NUP62, and β-actin, respectively. The ratio of these protein signals in cells transfected with RACK1 siRNA to those transfected with control siRNA is denoted as RACK1 (+:-), p-PKCα (+:-), p-PKCβ (+:-), p-NUP62 (+:-), and IBV N (+:-). For panel B and C, the value of mock-infected cells was regarded as 1.

### The pan-coronaviruses N proteins promotes the anchoring of p-PKCα to RACK1, and both PKCα/β and RACK1 are required for N protein to suppress the host antiviral response

One of the major functions of RACK1 is anchoring and trafficking activated PKC to specific subcellular locations [62]. We further investigated the effect of the interaction between IBV N and RACK1 on PKCα/β activity. Flag-tagged IBV N was expressed in DF-1 cells and immunoprecipitated with anti-Flag antibody, followed by Western blot analysis to detect endogenous RACK1, p-PKCα, and p-PKCβ. As depicted in Fig 11A, both endogenous RACK1 and p-PKCα were successfully co-immunoprecipitated with Flag-tagged IBV N, while p-PKCβ was not detected in the precipitated complex. This result demonstrates that IBV N, RACK1, and p-PKCα interact with each other to form a complex.

**Fig 11.**
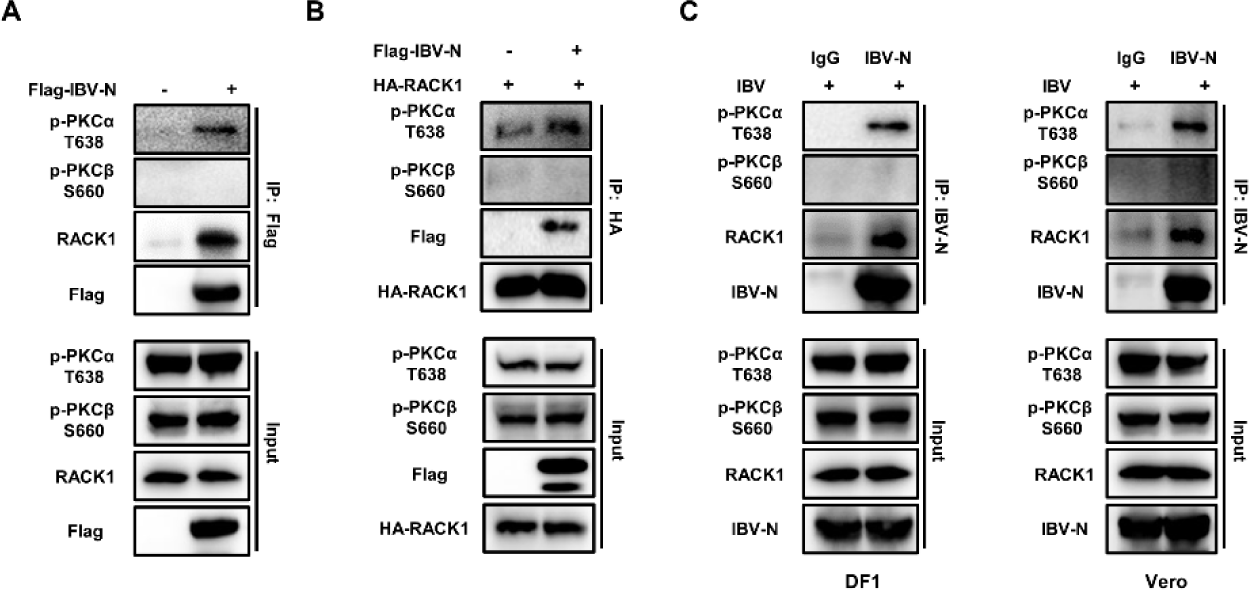
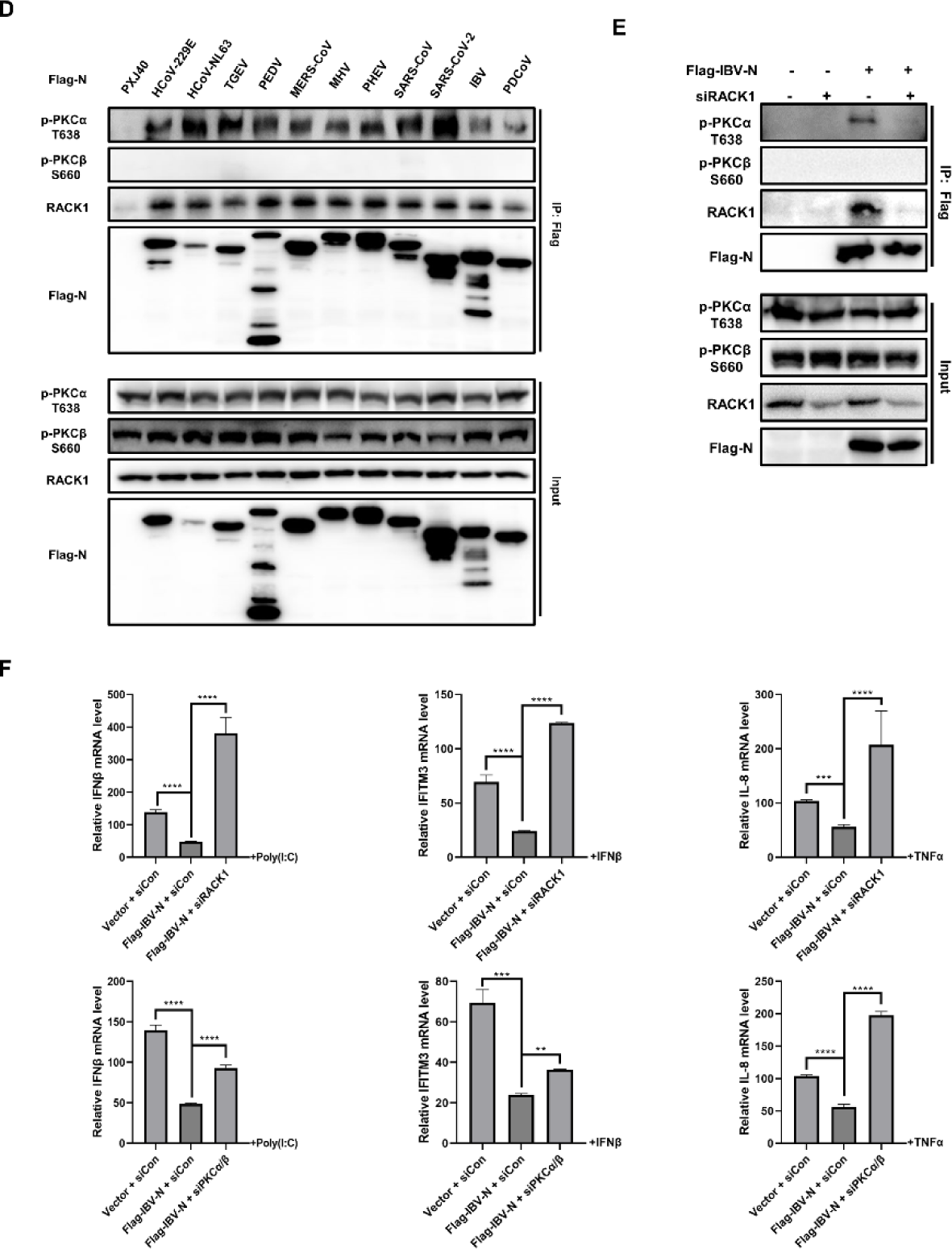
Interaction between IBV N, p-PKCα, and RACK1, and their role in suppression of innate immune response. (A) DF-1 cells were transfected with Flag-tagged IBV N or vector PXJ40 for 24 h and subjected to Co-IP using anti-Flag antibody, followed by Western blot analysis. (B) Plasmids encoding HA-RACK1 and Flag-tagged IBV N, or HA-RACK1 and PXJ40 were co-transfected into DF-1 cells for 24 h. Cell lysates were subjected to Co-IP using anti-HA antibody, followed by Western blot analysis. (C) DF-1 cells or Vero cells were infected with IBV for 12 h and subjected to Co-IP using anti-IBV N antibody. The interaction of IBV N, p-PKCα, p-PKCβ, and RACK1 was detected by Western blot. (D) HEK-293T cells were transfected with plasmids encoding Flag-tagged N proteins from eleven coronaviruses or PXJ40 for 24 h. Cell lysates were subjected to CO-IP with anti-Flag antibody, followed by immunoblot with indicated antibodies. (E) siRACK1 or control siRNA was transfected into DF-1 cells for 36 h, followed by transfection of plasmid encoding Flag-tagged IBV N or vector PXJ40 for 24 h. Cell lysates were subjected to Co-IP using anti-Flag antibody, followed by Western blot analysis. (F) siRNA targeting RACK1, PKCα/β, or control siRNA was transfected into DF-1 cells for 36 h, followed by transfection of plasmid encoding Flag-tagged IBV N or vector PXJ40. At 24 h post-transfection, cells were stimulated with poly(I:C), IFNβ, or TNFα, respectively. Cells were harvested after 12 h post-stimulation, and the transcription levels of IFNβ, IFITM3, and IL-8 was measured by qRT-PCR. The group transfected with vector PXJ40 without stimulation was set as 1.

In cells overexpressing HA-RACK1, more p-PKCα, but not p-PKCβ, was co-immunoprecipitated with HA-RACK1 in the presence of IBV N (Fig 11B), indicating that IBV N promotes the anchoring of p-PKCα to scaffold RACK1. This interaction was confirmed under IBV infection conditions (Fig 11C), as evidenced by successful pull-down of N protein together with endogenous RACK1 and p-PKCα using anti-IBV N polyclonal antibody, while p-PKCβ was not associated with RACK1 and N protein. These results validate the formation of a complex involving N, RACK1, and p-PKCα during IBV infection.

To determine whether the formation of the N-RACK1-p-PKCα complex is a conserved feature, Flag-tagged N proteins from eleven strains of coronaviruses were expressed in HEK-293T cells, and Co-IP was performed using anti-Flag antibody. HEK-293T cells were chosen for this study due to their high transfection efficiency, which facilitates the expression of N proteins. Western blot analysis showed that all the N proteins were able to bind and pull down endogenous RACK1 and p-PKCα together; once again, p-PKCβ was not detected in the precipitates. These results confirm that the formation of the N-RACK1-p-PKCα complex is conserved across pan-coronaviruses, and N protein plays an important role in promoting the anchoring of p-PKCα to RACK1 (Fig 11D).

Next, we investigated whether RACK1 serves as the scaffold to anchor N and PKCα together. RACK1 was knocked down in DF-1 cells, followed by transfection of IBV N. The Flag antibody successfully co-immunoprecipitated Flag-tagged IBV N and p-PKCα together with RACK1 in control siRNA-transfected cells. However, it failed to co-immunoprecipitate p-PKCα in the absence of RACK1 (siRACK-transfected cells). Once again, p-PKCβ was not co-immunoprecipitated with IBV N protein (Fig 11E). This result demonstrates that RACK1 serves as the scaffold for the formation of the N-RACK1-p-PKCα complex.

To test the effect of RACK1 and PKCα/β on the expression of antiviral genes, these proteins were individually knocked down in DF-1 cells, followed by the overexpression of IBV N and stimulation with poly(I:C), IFNβ, or TNFα. The transcription of IFNβ, IFITM3, and IL-8 was examined by qRT-PCR. As shown in Fig 11F, knockdown of either RACK1 or PKCα/β significantly restored the expression of these antiviral genes, which had been suppressed by IBV N. Therefore, both RACK1 and PKCα/β are required for N protein to suppress the innate immune response.

The anchoring of p-PKCα to the RACK1-N complex, rather than p-PKCβ, and the essential role of PKCα/β and RACK1 in NUP62 phosphorylation and N protein-mediated suppression of antiviral gene expression, suggest that the association with the RACK1-N complex enables p-PKCα to execute its kinase function in close proximity to its substrates, including phosphorylating NUP62. Conversely, since p-PKCβ does not associate with the RACK1-N complex, it likely exerts its kinase function by translocating to specific subcellular locations through alternative mechanisms.

### Nuclear export signal (NES) of IBV N is essential for promotion of cytoplasmic dispersion of FG-Nups and suppression of innate immune response

The coronavirus N protein compromises three highly conserved domains: the N-terminal viral RNA binding region (NTD), the Ser/Arg-rich region (SR-domain), and the C-terminal dimerization domain (CTD). To characterize which domain is involved inhibiting nucleocytoplasmic trafficking, plasmids encoding four IBV N truncation fragments were constructed: ΔNTD with deletion of N-terminal 1 to 160 aa, the viral RNA binding region; ΔSR with deletion of the Ser/Arg-rich region 165 to 190 aa, the potential phosphorylation region; ΔCTD with deletion of the C-terminal 215 to 409 aa, the dimerization domain; ΔNES by removing the nuclear export signal (NES, ^291^LQLDGLHL^298^) (Fig 12A). Vero cells were transfected with these plasmids and applied to immunofluorescence analysis. As illustrated in Fig 12B, both ΔNTD and ΔSR were distributed in the cytoplasm and exhibited the ability to inhibit the nuclear translocation of IRF3, p65, STAT1, and STAT2 in response to their respective stimuli, similar to wild-type IBV N. However, ΔCTD and ΔNES were primarily localized in the nucleus and showed a loss of capacity to inhibit the nuclear translocation of IRF3, p65, STAT1, and STAT2. The nuclear retention of ΔCTD might be attributed to the loss of NES (^291^LQLDGLHL^298^), which is located within the CTD (215 to 409 aa). Thus, the cytoplasmic distribution of N protein might be critical for perturbing the nucleocytoplasmic trafficking.

**Fig 12.**
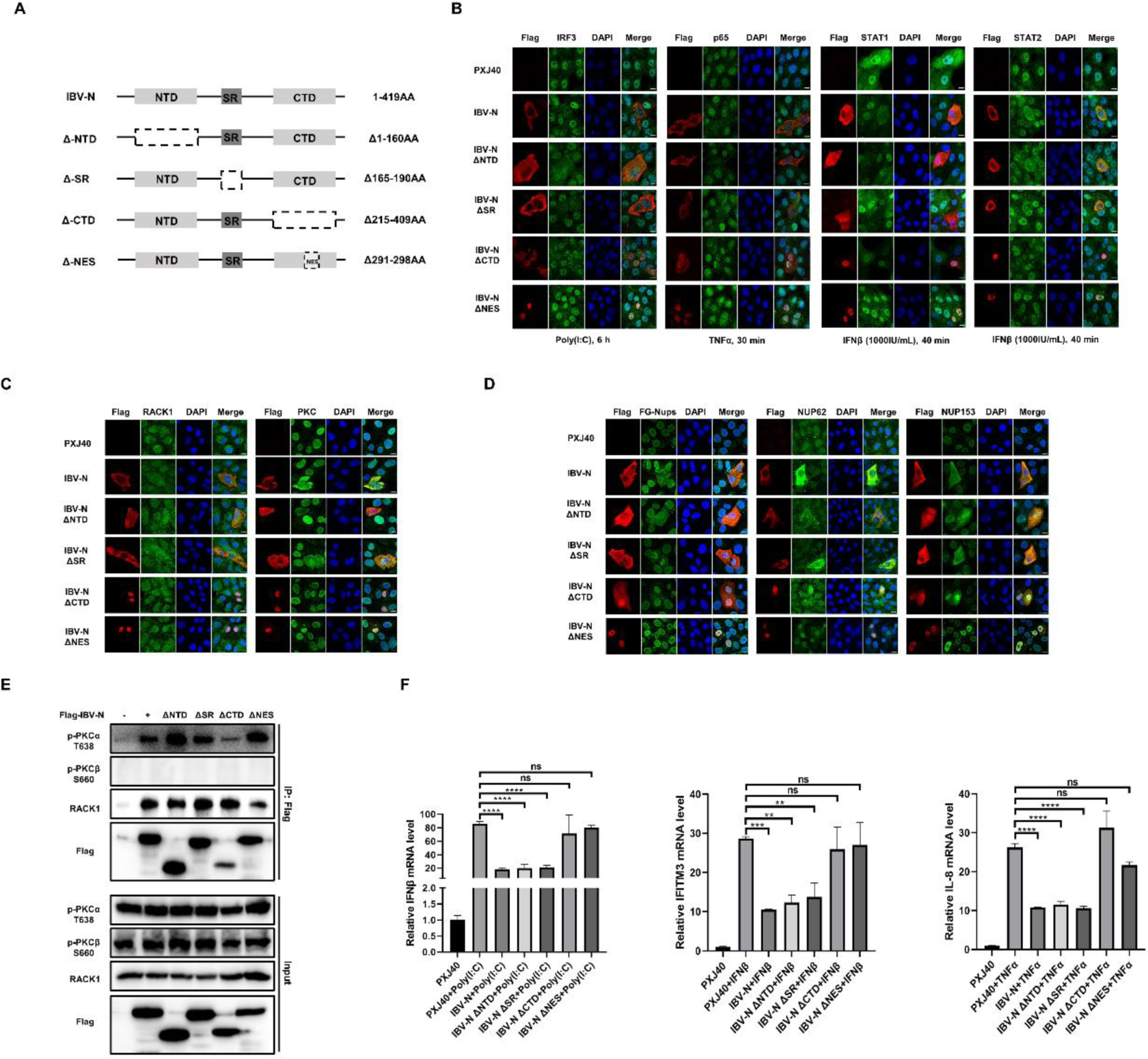
Significance of cytoplasmic localization of IBV N protein in disrupting the nuclear envelope localization of Nups and suppressing antiviral gene expression. (A) Schematic representation of truncated mutants of the IBV N protein. (B) Vero cells were transfected with plasmids encoding Flag-tagged IBV N, ΔNTD, ΔSR, ΔCTD, ΔNES, or PXJ40. At 18 h post-transfection, cells were treated with poly (I:C), while treatment with IFNβ or TNFα was initiated at 24 h post-transfection. Subsequently, cells were subjected to immunostaining. (C-D) Vero cells were transfected with plasmids encoding Flag-tagged IBV N, ΔNTD, ΔSR, ΔCTD, ΔNES, or PXJ40. At 24 h post-transfection, cells were subjected to immunostaining with corresponding antibodies. Representative images from three independent experiments are shown. Scale bars: 10 μm. (E) DF-1 cells were transfected with plasmids encoding Flag-tagged IBV N or the truncated mutants, or PXJ40. At 24 h post-transfection, cell lysates were subjected to Co-IP with anti-Flag antibody and subsequently immunoblotted with corresponding antibodies. (F) DF-1 cells were transfected with plasmids encoding Flag-tagged IBV N, the truncated mutants, or PXJ40. At 24 h post-transfection, cells were treated with poly(I:C), IFNβ, or TNFα for 12 h. Cells were harvested, and the levels of IFNβ, IFITM3, and IL-8 were determined using qRT-PCR.

We further investigated the subcellular localization of RACK1 and PKCα/β in cells expressing various IBV N mutants. As illustrated in Fig 12C, cells transfected with PXJ40 exhibited RACK1 signal in the nucleus and perinuclear region, whereas PKCα/β signals were predominantly detected in the nucleus. In cells expressing wild-type IBV N, ΔNTD, or ΔSR, RACK1 and PKCα/β were redistributed to the cytoplasm, colocalizing with N protein or its truncated mutants. This indicates that N protein has the capacity to modulate the subcellular localization of RACK1 and PKCα/β. However, when ΔCTD and ΔNES mutants were confined to the nucleus, there was no significant dispersion of RACK1 and PKCα/β into the cytoplasm: RACK1 remained localized in the nucleus and perinuclear region, while PKCα/β predominantly remained in the nucleus (Fig 12C). These findings suggest that the localization of the N protein dictates the positioning of the RACK1-PKCα complex. Further examination of the impact of these N mutants on the intracellular distribution of Nups revealed that ΔNTD and ΔSR induced cytoplasmic dispersion of FG-Nups, NUP62, and NUP153, similar to cells expressing wild-type N protein. Conversely, in cells expressing ΔNES, FG-Nups, NUP62, and NUP153 signals remained concentrated at the NE, with intense signals observed within the nucleus (Fig 12D). Notably, in cells expressing ΔCTD, a minor fraction of FG-Nups and NUP153 exhibited cytoplasmic dispersion, while the signal of NUP62 and NUP153 was intensified in the nucleus. Co-IP analysis revealed that similar to wild-type N protein, ΔNTD, ΔSR, ΔCTD, and ΔNES exhibited varying degrees of interaction capability with RACK1 and p-PKCα (Fig 12E).

Consistent with the observations in Fig 12B-D, ΔNTD and ΔSR maintained the capability to inhibit the expression of IFNβ, IFITM3, and IL-8 in response to poly(I:C), IFNβ, and TNFα stimuli, whereas ΔCTD and ΔNES lost this capacity (Fig 12F). Overall, the presence of CTD and NES enables the N protein to localize in the cytosol alongside RACK1 and PKCα/β, thereby facilitating the function of the N-RACK1-p-PKCα complex in inducing cytoplasmic dispersion of FG-Nups. This process prevents the nuclear translocation of transcription factors and the subsequent antiviral innate immune response, as indicated by the results presented in Fig 12B-F.

## Discussion

The coronavirus has evolved multiple strategies to inhibit the innate immune response for its own benefit. In this study, we demonstrate that IBV infection inhibits the expression of several antiviral genes by suppressing the nuclear translocation of their corresponding transcription factors: IRF3, STAT1/STAT2/IRF9, and p65. We identified the IBV N protein as the factor responsible for retaining these transcription factors in the cytoplasm and suppressing the expression of antiviral genes. Both IBV infection and N protein expression promote the cytoplasmic dispersion of multiple Nups, indicating perturbation of the NPC function, which governs the nucleocytoplasmic trafficking of RNA and proteins. Although immunofluorescence analysis shows the colocalization of N and Nups in the cytoplasm, there is no direct interaction between the N protein and several Nups (NUP62 and NUP42), as determined by Co-IP analysis (S5 Fig). Previous studies have reported that the disassembly of the NPC is regulated by phosphorylation of Nups during cell mitosis [48, 63]. Here, Western blot analysis demonstrates that NUP62 is phosphorylated at T269 and S272 during IBV infection. Chemical inhibition of intracellular PP1 and PP2A activity by okadaic acid leads to the accumulation of phosphorylated NUP62 and promotes its cytoplasmic dispersion. These findings strongly support the idea that phosphorylation events of Nups lead to the disassembly of the NPC. Mechanistic studies reveal that the N protein interacts with RACK1 and promotes the anchoring of p-PKCα, but not p-PKCβ, to RACK1. The presence of both RACK1 and PKCα/β is required for the phosphorylation of NUP62 and suppression of antiviral gene expression, thereby benefiting virus infection. Inhibition of PKCα/β activity by chemical inhibitor prevents the cytoplasmic distribution of multiple Nups in both N protein-expressing or IBV-infected cells, further indicating the role of PKCα/β in Nups phosphorylation and NPC disassembly. Although phosphorylation of other Nups was not detected due to the lack of phospho-specific antibodies, the cytoplasmic dispersion of FG-Nups, NUP42, NUP153, and TPR suggests that additional Nups might undergo phosphorylation during IBV infection. These observations were also apparent in cells expressing the N proteins across various coronaviruses. Hence, coronavirus infection potentially triggers the phosphorylation of Nups and the disassembly of the NPC through the N-RACK1-p-PKCα-Nup signaling axis. Disruption of the NPC hinders transcription factors from accessing the nucleus, consequently inhibiting the transcription of antiviral genes and ultimately leading to immune suppression, thereby favoring viral replication (Fig 13).

**Fig 13.**
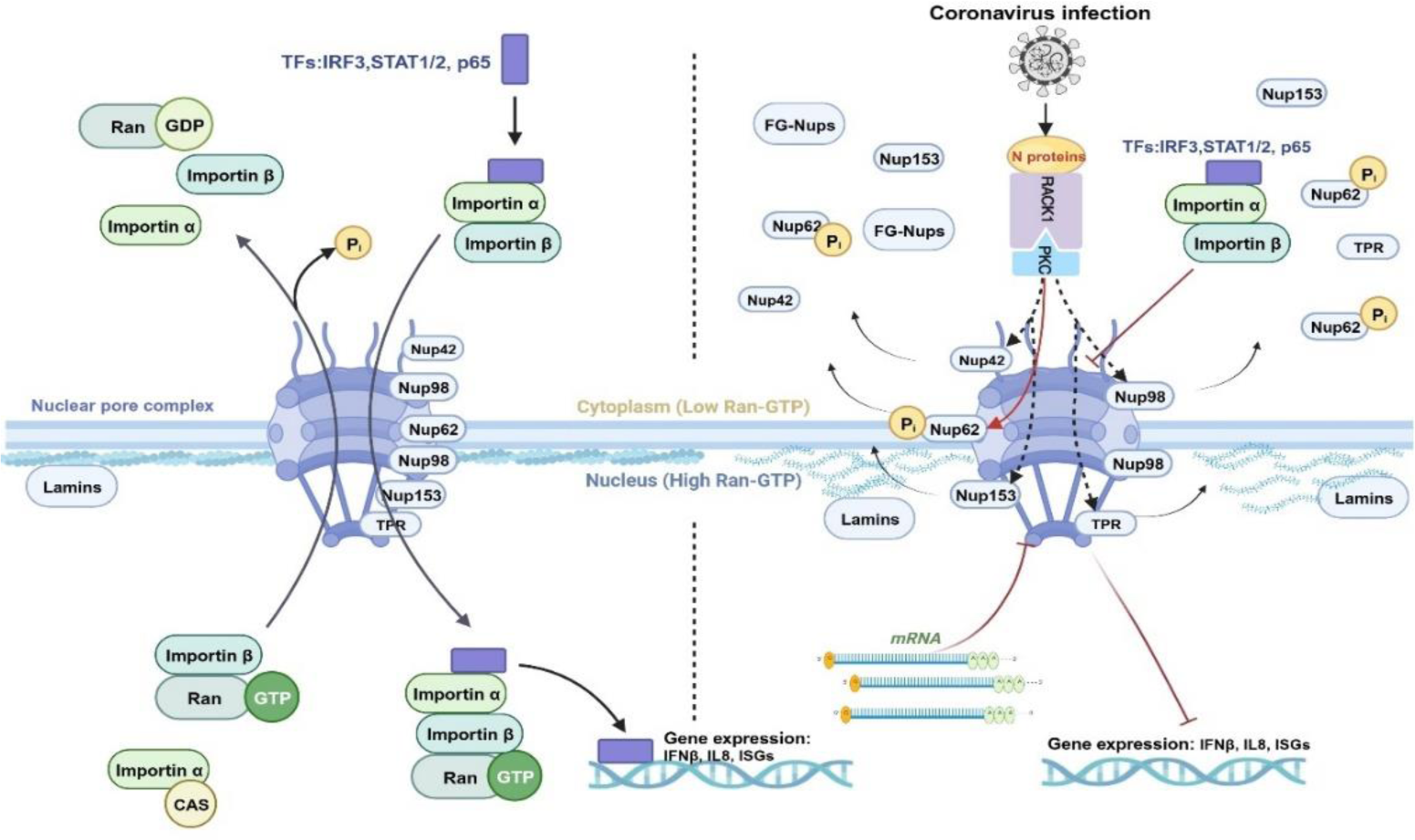
A working model illustrating how coronavirus N protein disrupts the nuclear transport system to inhibit the nuclear translocation of transcription factors and subsequent gene expression. In the left panel, the canonical nuclear import pathway is described, where the nuclear localization signal (NLS) of cargo proteins (e.g., transcription factors) is recognized by nuclear import receptors such as importin α, forming a complex with importin β. This complex then translocates through NPC by interacting with FG-Nups. Upon entering the nucleoplasm, importin β binds Ran-GTP, leading to disassembly of the import complex and release of the cargo. Importin β bound to Ran-GTP is transported back to the cytoplasm, while importin α is recycled by cellular apoptosis susceptibility (CAS) protein (also known as exportin 2). GTP hydrolysis of Ran releases importin β for the next round of import.

**S5 Fig.**
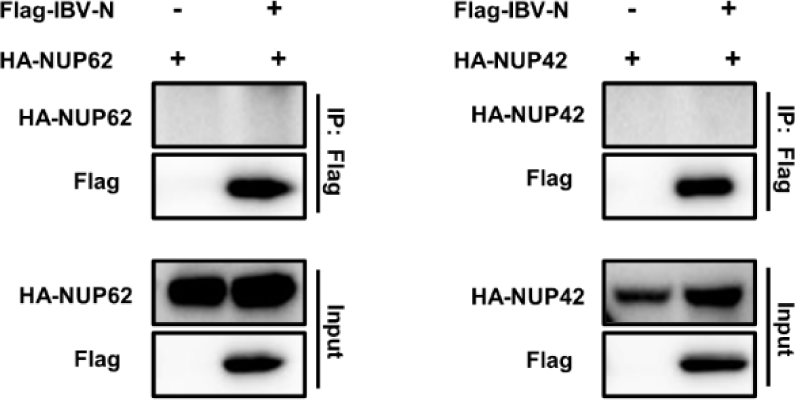
The in vitro interaction results between IBV N protein and NUP62 or NUP42. Plasmids encoding HA-NUP62 or HA-NUP42 were co-transfected with Flag-tagged IBV N protein, or PXJ40 control plasmid, into HEK-293T cells for 24 h. Cell lysates were subjected to Co-IP using anti-Flag antibody, followed by Western blot analysis.

In the right panel, during coronavirus infection, the N protein interacts with RACK1 and recruits p-PKCα to RACK1 to form a ternary complex. This N-RACK1-p-PKCα complex phosphorylates NUP62, leading to its cytoplasmic dispersion together with other Nups. Consequently, the NPC fails to transport cargo into the nucleus, thereby inhibiting the nuclear translocation of transcription factors such as IRF3, STAT1/2, and p65, subsequently blocking the expression of downstream antiviral genes. This figure was created using Biorender (BioRender, Toronto, ON, Canada).

FG-Nups primarily anchor to the central channel of the NPC, forming a dense barrier that prevents passive diffusion while facilitating the passage of cargos via nuclear transport receptors [64]. During IBV infection, no degradation of FG-Nups is observed; however, NUP62 and NUP42 exhibit a shift in size, and phosphorylation of NUP62 at T269 and S272 is confirmed by anti-p-NUP62 antibody (Fig 2D). In Vero cells, a potential cleaved p-NUP62 band at around 45 kDa is detected, indicating the instability of p-NUP62 during infection. Extensive phosphorylation of Nups has been reported to disrupt protein-protein interactions at key contact nodes within the NPC, leading to NPC disintegration and dispersion of Nups into the cytosol [48]. Although we were unable to detect phosphorylation of NUP98 by Western blot analysis due to the lack of an anti-p-NUP98 antibody, immunofluorescence analysis revealed the breakdown of the NUP98 nuclear ring signal in cells expressing N proteins, indicating disassembly of this Nup (Fig 5A). The central FG-Nup subunit NUP62 is demonstrated to be phosphorylated and disperses into the cytosol, along with the cytoplasmic dispersion of Nup153, Nup42, and TPR (Fig 2A, 2C). When the PP1 and PP2A inhibitor okadaic acid was applied to cells, the phosphorylation level of NUP62 was dramatically increased (Fig S4A), which coincided with cytoplasmic dispersion (Fig S4B). This observation demonstrates a direct correlation between the phosphorylation of NUP62 and its diffusion into the cytosol. It is noteworthy that in cells treated with okadaic acid, the cytoplasmic distribution pattern of FG-Nups (Fig S4B) is not as pronounced as in IBV-infected or N protein-expressing cells; most FG-Nups remain associated with the NE. Thus, in virus-infected cells or cells expressing the N protein, the NPC is dismantled more extensively than in cells treated with a PP1 and PP2A inhibitor. Moreover, inhibition of phosphatase activity by inhibitor results in the appearance of two p-NUP62 bands around 120 kDa and 62 kDa (Fig S4A), indicating the formation of dimers after hyperphosphorylation. The antibody against total NUP62 detects three bands adjacent to 62 kDa in inhibitor-treated cells (Fig S4A), suggesting that NUP62 harbors multiple phosphorylation sites in addition to T269 and S272. It has been reported that extracellular signal-regulated kinase (ERK) and p38 MAPK, activated by encephalomyocarditis virus leader protein, are involved in hyperphosphorylation of several FG-Nups including NUP62, NUP153, and NUP214, leading to diffusion of their nuclear ring signals and inhibition of nuclear import [65, 66]. The ERK-targeted phosphorylation site of NUP62 has previously been mapped to a single PxTP motif within the FG repeat region of NUP62, resulting in an alteration in NPC sensitivity to STAT3 passage [67]. In our study, the phosphorylation site of NUP62 was identified at T269 and S272 within the flexible region. Hyperphosphorylation of NUP62 coincided with cytoplasmic dispersion of NUP62 itself and multiple other Nups, as well as inhibition of transcription factor import. Thus, we conclude that the perturbation of NPC integrity in IBV-infected cells is attributed to the phosphorylation events of NUP62 and other FG-Nups.

It has been elucidated that the disintegration of the nuclear envelope induced by parvovirus infection involves a sequential enzymatic cascade mediated by PKC, CDK2, and caspase-3 [68]. In this study, we focused on screening the kinases responsible for NUP62 phosphorylation using chemical inhibitors, which prompted us to investigate PKCα/β. PKCα/β is known for its ability to phosphorylate a diverse set of protein substrates and its involvement in various cellular processes, including cell adhesion, cell transformation, cell cycle regulation, apoptosis, and macrophage development [69]. Several direct substrates of PKCα/β have been identified, such as RAF1 [70], BCL2 [71], DOCK8 [72], Lamin B1 [55] and components of the signaling cascade involving ERK1/2 [73], as well as RAP1GAP [74]. In our study, we observed that the PKCα/β specific inhibitor, Enzastaurin, effectively suppresses the cytoplasmic dispersion of several Nups in both IBV-infected cells and cells expressing the N proteins of pan-coronaviruses (Fig 8A and 9C), indicating a clear correlation between PKCα/β activity and Nups disassembly. Inhibition of PKCα/β activity or knockdown of PKCα/β specifically reduces the level of phosphorylated NUP62 (Fig 9D, 9E). This study provides the first evidence of PKCα/β involvement in NUP62 phosphorylation and subsequent cytoplasmic dispersion. Additionally, the presence of PKCα/β is essential for IBV or N protein-mediated suppression of antiviral gene expression (Fig 9F and Fig 11F) and facilitates IBV infection (Fig 9 D-E). In line with our findings, a recent study has demonstrated that the replication of SARS-CoV-2 is impeded by pan-PKC inhibitors such as Go 6983, Bisindolmaleimide I, Enzastaurin, and Sotrastaurin [75]. This suggests that PKC may play an essential role in facilitating coronavirus infection. Hence, the targeting of PKC emerges as a promising strategy in the development of broad-spectrum anti-coronaviral drugs.

The regulation of PKC signaling by coronavirus involves the interaction of the viral N protein with the PKC scaffold protein RACK1, as revealed by N protein interactome analysis and Co-IP. As a highly conserved multifunctional protein, RACK1 interacts directly or in complex with various cellular proteins, including PKCα/βⅡ, contributing to protein shuttling, subcellular localization, and activity modulation [54, 76]. In IBV-infected or pan-coronaviruses N protein-expressing cells, we observed an augmentation in the interaction between p-PKCα and RACK1 (Fig 11A-C), with PKCα/β and RACK1 exhibiting colocalization with the N protein in the cytoplasm (Fig 7C and 9B). These findings suggest that the N protein facilitates the translocation of p-PKCα from the nucleus to the cytoplasm, where it associates with RACK1. The depletion of RACK1 demonstrates its indispensability in the formation of the N-RACK1-p-PKCα complex (Fig 11E). Furthermore, the presence of RACK1 is essential for the phosphorylation of PKCα, PKCβ, and NUP62, as well as for efficient IBV replication (Fig 10A). The conserved enhancement of p-PKCα anchoring to RACK1 by the N protein across pan-coronaviruses suggests a common mechanism employed by coronaviruses to recruit p-PKCα to RACK1, promoting the phosphorylation and disassembly of NUP62 (Fig 11D), as well as potentially affecting other Nups. The necessity of RACK1 and PKCα/β for the N proteins or IBV to suppress the expression of antiviral factors such as IFNβ, IFITM3, and IL-8 (Fig 9F, 10C, and 11F) further underscores the critical role of the N-RACK1-p-PKCα complex in antagonizing the host innate immune response, potentially through phosphorylation events on NUP62 or other substrates. In line with our findings, RACK1 has been implicated in facilitating SARS-CoV-2 replication, as its depletion has been shown to reduce infectious virus release and intracellular spike protein expression [77]. Furthermore, it has been demonstrated that the N protein of porcine reproductive and respiratory syndrome virus (PRRSV) interacts with RACK1, thereby facilitating PRRSV replication [78]. Thus, RACK1 may play a pivotal role not only in coronavirus infections but also in arterivirus infections. Previous research indicates that a portion of the RACK1-binding site for PKCβII resides within the PKCβII V5 domain, and a peptide corresponding to amino acids 645-650 in PKCβII selectively inhibits phorbol 12-myristate 13-acetate (PMA)-induced translocation of PKCβII, thereby blocking PKC activity [79]. Developing RACK-competitive PKC inhibitors could be a novel strategy for the development of anti-coronaviral therapeutics.

As a multifunctional protein, the coronavirus N protein plays pivotal roles in packaging viral RNA into ribonucleoprotein, participating in virion assembly, modulating viral replication and transcription, and regulating host innate immunity [80, 81]. Our study reveals that pan-coronaviruses N proteins interact with RACK1 and p-PKCα (Fig 11D), as well as induce the cytoplasmic dispersion of multiple Nups (Fig 5A), leading to the blockade of nuclear translocation of transcription factors such as IRF3, STAT1/STAT2/IRF9, and p65 (Fig 5C-5D), ultimately inhibiting the expression of antiviral genes (Fig 5E). The N protein consist of three conserved domains: NTD, SR domain, and CTD [82–84]. Using IBV N protein as a model, we identified the NES residing in NTD as the primary sequence responsible for the cytoplasmic distribution of the N-RACK1-p-PKCα complex and Nups, prevention of the nuclear import of transcription factors, and repression of antiviral gene expression (Fig 12). Our findings suggest that within the ternary complex of IBV N, RACK1, and p-PKCα, N plays a pivotal role in determining the subcellular localization of the complex. The full-length N protein, ΔNTD, ΔSR, interact with RACK1 and p-PKCα, causing their distribution in the cytoplasm. However, N protein lacking CTD or NES is retained in the nucleus together with RACK1 and PKCα/β, thereby losing the ability to promote the cytoplasmic dispersion of Nups and subsequent nuclear importing and transcription events. Thus, the cytoplasmic localization of IBV N determines the positioning and regulation role of the N-RACK1-p-PKCα complex in phosphorylating NUP62 or other substrates. We attempted to generate a recombinant virus by deleting the NES of N protein based on the IBV Beaudette strain using reverse genetic technique; however, we were unable to rescue the NES-deficient rIBV strain. This underscores the significance of nuclear export and cytoplasmic localization for the N protein to fulfill its function, which is indispensable for IBV replication. Although limited research has been conducted on the NES of N proteins from other coronaviruses, localization to the nucleolus appears to be a common feature of N proteins from four genera of coronaviruses [85, 86]. This suggests that the N protein possesses the capability for cytoplasm-nucleus shuttling; for instance, phosphorylated SARS-CoV N is translocated to the cytoplasm from nucleus with the assistance of 14-3-3 [87]. Given the critical role of N protein subcellular localization in its function, investigating the nuclear import and export strategies of coronavirus N proteins will presents an intriguing avenue for future research.

The disassembly of NPC not only inhibits the nuclear transport of transcription factors involved in the immune response but also disrupts host mRNA export or sequesters nuclear proteins essential for viral replication in the cytoplasm, thereby promoting viral replication. In addition to inhibiting the nuclear translocation of STAT1, a recent study demonstrated that SARS-CoV and SARS-CoV-2 ORF6 interacts with NUP98/Rae1 to impede cellular mRNA export, thereby reducing the translation of antiviral genes and diverting limited cellular translational machinery towards viral translation [88]. Future investigations are warranted to elucidate whether N protein prevents host mRNA export to reduce host translation events or retains nuclear proteins in the cytoplasm to facilitate virus replication.

In summary, our study reveals that IBV N protein promotes the anchoring of p-PKCα to RACK1 and relocates the N-RACK1-p-PKCα complex to the cytoplasm. In this context, p-PKCα phosphorylates NUP62 and potentially other Nups, promoting their disassembly and cytoplasmic dispersion. Consequently, this process inhibits the nuclear import of transcription factors such as IRF3, STAT1/2/IRF9, and p65, thereby blocking the expression of antiviral genes, and ultimately facilitating IBV replication. The disruption of nuclear trafficking and inhibition of transcription factor nuclear entry represent a novel and evolutionarily conserved function in N proteins across pan-coronaviruses.

## Materials and methods

### Cells and viruses

Chicken embryo fibroblast DF-1 cells (ATCC® CRL-12203™), African green monkey kidney epithelial Vero cells (ATCC® CCL-81™), and human embryonic kidney HEK-293T cells (ATCC® CRL-3216™) were obtained from ATCC. These cells were cultured in Dulbecco’s Modified Eagle Medium (DMEM) supplemented with 10% (v/v) fetal bovine serum (FBS) (Gibco-Thermo Fisher, Waltham, MA, USA). The IBV Beaudette strain was kindly provided by Prof. Dingxiang Liu’s laboratory at South China Agricultural University.

### Antibodies and chemicals

Rabbit anti-IBV-N and mouse anti-IBV-N polyclonal antibodies were generated in our laboratory. Additionally, the following antibodies were purchased: rabbit anti-IRF3 (ab68481), rabbit anti-p65 (ab32536), rabbit anti-p38 (ab170099), rabbit anti-IRF-9 (ab271043), rabbit anti-TPR (ab170940), rabbit anti-Ran (ab157213), rabbit anti-NUP42 (Nucleoporin hCG1, ab192609), rat anti-NUP62 (ab188413), mouse anti-NUP153 (ab24700), mouse anti-FG-Nups [Mab414] (ab24609), mouse anti-importin β1 (ab2811), rabbit anti-PKCα/βⅡ (ab184746), rabbit anti-phospho-PKCα (T638) (ab32502), rabbit anti-NUP98 (#2598), rabbit anti-phospho-PKCβII (Ser660) (#9371), rabbit anti-STAT1 (#14994), rabbit anti-HA Tag (#3724), rabbit anti-Flag Tag (#14793), rabbit anti-PP1α (#2582), rabbit anti-PP2A C Subunit (#2038), rabbit anti-STAT2 (16674-1-AP), rabbit anti-NUP62 (13916-1-AP), rabbit anti-importin α1 (16674-1-AP), and rabbit anti-RACK1 (R1905) were purchased from Abcam, Cell Signaling Technology, Proteintech Group, and Merck, respectively. Mouse anti-Flag Tag (M185-3L) was purchased from MBL, and rabbit anti-β-actin (AC026) was also used. The dilution of antibodies and their cross-reactivity with corresponding chicken proteins are summarized in Table 1. Alexa Fluor goat anti-rabbit-488 (A-11034), Alexa Fluor goat anti-rabbit-594 (A-11037), Alexa Fluor goat anti-mouse-488 (A-11029), and Alexa Fluor goat anti-mouse-594 (A-11005) were obtained from Invitrogen, USA. Poly(I:C) (31852-29-6) was from InvivoGen, France. Recombinant human IFN-β protein (#8499-IF) was purchased from Bio-Techne R&D Systems, USA. Recombinant Human TNF-α (P00029) was purchased from Solarbio, China. Okadaic Acid (#5934) was purchased from Cell Signaling Technology, USA. Enzastaurin (HY-10342) was purchased from MCE, China. The ClonExpress Ultra One Step Cloning Kit (C115) and Mut Express II Fast Mutagenesis Kit V2 (C214) were purchased from Vazyme, China.

**Table 1.** Dilution of primary antibodies and their cross-reactivity with corresponding chicken proteins.

| Antibody name | Application and dilution | Cross-reactivity with chicken protein |
| --- | --- | --- |
| Anti-IBV-N | Western blot (1:5000)<br>Immunofluorescence (1:500) | NA<br>NA |
| Anti-IRF3 | Immunofluorescence (1:100) | Yes |
| Anti-p65 | Immunofluorescence (1:100) | Yes |
| Anti-STAT1 | Immunofluorescence (1:100) | No |
| Anti-STAT2 | Immunofluorescence (1:100) | Yes |
| Anti-IRF9 | Immunofluorescence (1:100) | No |
| Anti-p38 | Immunofluorescence (1:100) | No |
| Anti-NUP62 | Western blot (1:1000)<br>Immunofluorescence (1:100) | Yes<br>Yes |
| Anti-p-NUP62 | Western blot (1:1000) | Yes |
| Anti-FG-Nups (mAb414) | Immunofluorescence (1:500) | No |
| Anti-NUP153 | Western blot (1:1000)<br>Immunofluorescence (1:500) | Yes<br>Yes |
| Anti-NUP98 | Western blot (1:1000)<br>Immunofluorescence (1:100) | Yes<br>Yes |
| Anti-NUP42 (hCG1) | Western blot (1:1000)<br>Immunofluorescence (1:50) | Yes<br>No |
| Anti-TPR | Western blot (1:1000)<br>Immunofluorescence (1:400) | Yes<br>Yes |
| Anti-Ran | Western blot (1:1000)<br>Immunofluorescence (1:100) | Yes<br>Yes |
| Anti-Importin β1 | Western blot (1:1000)<br>Immunofluorescence (1:100) | Yes<br>No |
| Anti-Importin α1 | Western blot (1:1000)<br>Immunofluorescence (1:100) | Yes<br>No |
| Anti-RACK1 | Western blot (1:1000)<br>Immunofluorescence (1:100) | Yes<br>No |
| Anti-PKC$\alpha/\beta$ | Western blot (1:1000)<br>Immunofluorescence (1:200) | Yes<br>Yes |
| Anti-p-PKCα | Western blot (1:1000) | Yes |
| Anti-p-PKCβ | Western blot (1:1000) | Yes |
| Anti-HA | Western blot (1:1000)<br>Immunofluorescence (1:500) | NA<br>NA |
| Anti-Flag | Western blot (1:1000)<br>Immunofluorescence (1:500) | NA<br>NA |
| Anti-PP1α | Western blot (1:1000) | Yes |
| Anti-PP2A C | Western blot (1:1000) | Yes |
| Anti-β-actin | Western blot (1:1000) | Yes |

### Plasmids construction

Plasmids encoding HCOV-229E-N, HCOV-NL63-N, TGEV-N, PEDV-N, MERS-CoV-N, MHV-N, PHEV-N, SARS-CoV-N, and SARS-CoV-2-N were provided by Prof. Tongling Shan (Shanghai Veterinary Research Institute, CAAS) [89]. Plasmids encoding Flag-tagged IBV nsp2, nsp3, nsp5, nsp6, nsp7, nsp8, nsp9, nsp12, nsp13, nsp14, nsp15, nsp16, E, M, 5a, and N, constructed by Dr. Gao, are maintained in our laboratory [90]. The plasmid encoding IBV N was generated by amplifying cDNA from IBV Beaudette-infected DF-1 cells using corresponding primers and cloning into PXJ40. IBV N ΔNTD, ΔSR, ΔCTD, and ΔNES mutants were generated by mutagenesis of the Flag-tagged IBV N plasmid using the Mut Express II Fast Mutagenesis Kit V2. PKCα and PKCβ genes were synthesized and ligated into the pCMV-HA expression vector by Sangon Bioengineering (Shanghai) Co., Ltd., Shanghai, China. NUP62, NUP42, and RACK1 genes were cloned by RT-PCR from HEK-293T cells and ligated into the pCMV-HA expression vector using the ClonExpress Ultra One Step Cloning Kit. The corresponding primers used to generate the above plasmids are shown in Table 2.

**Table 2.**
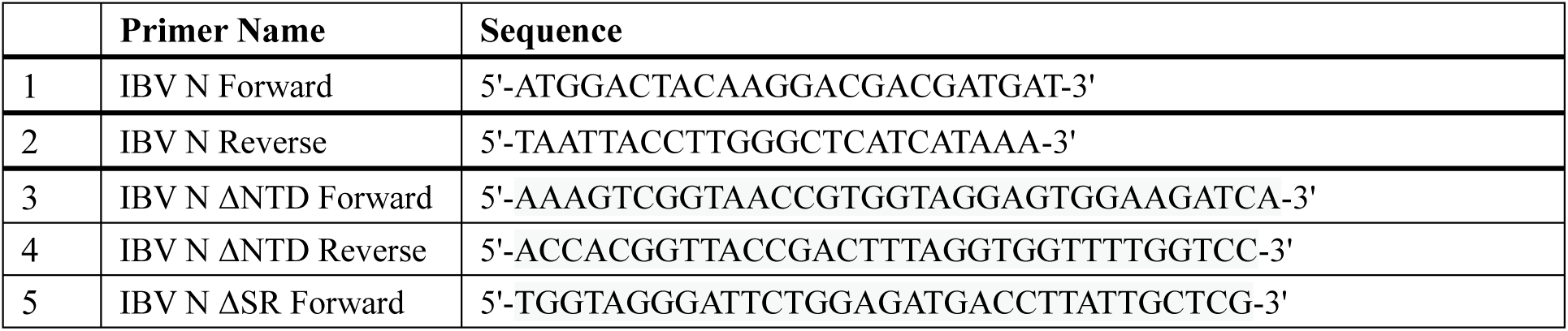
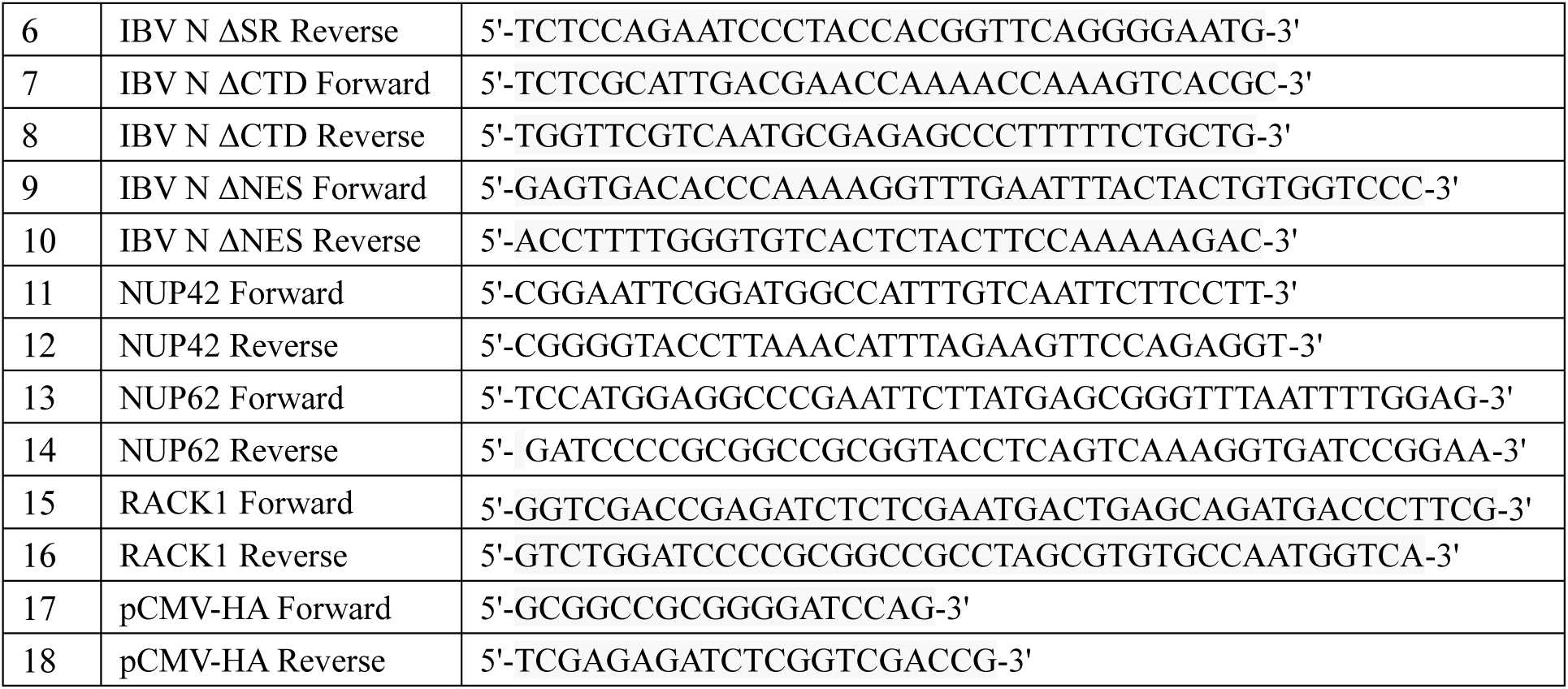
Primer sequences used for plasmid construction.

### Cell transfection and RNA interference

Vero cells or DF-1 cells were seeded in 6-well plates, 12-well plates, or chamber slides (Nunc Lab-Tek II Chamber Slide System, Thermo Fisher Scientific, USA) with 70-80% confluency. The indicated plasmids were transfected into cells using Lipofectamine 2000 (Invitrogen, Carlsbad, CA) according to the manufacturer’s instructions. Briefly, 1 μg of plasmid and 3 μL of Lipofectamine 2000 (m/v = 1:3) were diluted in 0.1 mL of Opti-MEM (Gibco, 31985070, Gaithersburg, MD). After 5 min of incubation, the plasmid and Lipofectamine 2000 were mixed and incubated at room temperature for 20 min to allow the formation of lipid-plasmid complexes. Finally, the complexes were added to the cultured cells and incubated for 24 h.

To knock down chicken RACK1 and PKCα/β genes in DF-1 cells, siGenome Gallus gallus RACK1 and PKCα/β siRNA were purchased from GenePharma Co, China. The sequences targeting RACK1 and PKCα/β were as follows: RACK1-siRNA: 5’-CGGGAUAUCUGAACACAGUTT-3’; PKCα/β-siRNA: 5’-GGAGCUCUAUGCAAUCAAATT-3’. A non-targeting control siRNA was also provided by GenePharma Co and used as a control with no specific gene targeting. For each siRNA transfection, 100 pmol of siRNA and 5 μL of Lipofectamine 2000 were diluted in 0.1 mL of Opti-MEM, respectively. After 5 min of incubation, the siRNA and Lipofectamine 2000 were mixed and incubated at room temperature for 20 min, allowing the formation of lipid-siRNA complexes. The complexes were then added to the cultured cells (30-40% confluency) and incubated for 48 h, followed by plasmid transfection or IBV infection. Cells were subjected to Western blot analysis or real-time qRT-PCR analysis at the indicated times.

### Western blotting analysis

DF-1 cells or Vero cells were seeded in 6-well plates and transfected with various siRNAs or infected with IBV at an MOI of 1 according to experimental requirements. Cells were harvested at the indicated time points or treated with Enzastaurin (1μM, 16 h) or Okadaic Acid (1μM, 1 h), with DMSO included in the parallel experiment as a negative control. Cell samples were lysed in 2x protein loading buffer (20 mM Tris-HCl, 2% SDS, 100 mM DTT, 20% glycerol, 0.016% bromophenol blue) and incubated in a 100°C metal bath for 10 min to fully denature the proteins. The denatured cell samples were then subjected to centrifugation at 12,000 rpm for 5 min. The supernatant proteins were resolved on a 10% SDS-PAGE and transferred to a nitrocellulose membrane (0.45 μm, Millipore, USA). Membranes were blocked in blocking buffer (5% nonfat milk, TBS, 0.1% Tween 20) for 1 h, followed by overnight incubation at 4°C with primary antibodies diluted in dilution buffer (Beyotime, P0023, China) as indicated in Table 1. The membranes were then incubated with secondary antibodies conjugated with HRP (Invitrogen, USA) diluted 1:10,000 in blocking buffer for 1 h at room temperature. After each incubation, membranes were washed three times with washing buffer (0.1% Tween in TBS). Proteins were visualized using the ECL detection system (Thermo, Rockford, IL). Image J program (NIH, USA) was used to quantify the intensities of corresponding bands on the Western blot according to the manufacturer’s instructions.

### Indirect immunofluorescence analysis

Cells were seeded onto chamber slides and transfected with various plasmids or infected with IBV at an MOI of 1, according to experimental requirements. At the indicated time points, cells were transfected with poly (I:C) (20 μg/mL) or treated with IFNβ (1000 IU/mL, 40 min), TNFα (20 ng/mL, 30 min), UV irradiation (1.92 J/cm2, 20 min), Enzastaurin (1 μM, 1 h), or Okadaic Acid (1 μM, 1 h). DMSO was used as the negative control in the case of drug treatment. Following treatment, cells were fixed with 4% paraformaldehyde for 15 min at room temperature. After three washes with PBS, cells were permeabilized with 0.5% Triton X-100 for 15 min and incubated in blocking buffer (3% BSA in PBS) for 1 h. Cells were then incubated with the primary antibody diluted in blocking buffer for 2 h at 37°C (the dilution was indicated in Table 1), followed by incubation with Alexa Fluor-conjugated secondary antibody diluted 1:500 in blocking buffer for 1 h at 37°C. In the case of double staining, cells were further incubated with the other primary antibody, followed by incubation with the corresponding fluorescent-conjugated secondary antibody. After each incubation step, the cells were washed three times with PBST. DAPI (Beyotime, C1002, China) was then applied to stain the nuclei for 10 min. Finally, cells were washed three times with PBST, and the subcellular localization of corresponding proteins was examined using a Zeiss LSM880 confocal microscope.

### Real-time quantitative RT-PCR analysis

DF-1 cells or HEK-293T cells were seeded in 6-well plates and transfected with siRNA or plasmid, or infected with IBV at the indicated MOI according to experimental requirements. At the specified time points, cells were transfected with poly I:C (20 μg/mL) or treated with IFNβ (1000 IU/mL) or TNFα (20 ng/mL). DMSO treatment was included in parallel experiments as a negative control.

Total cellular RNA was extracted using Trizol reagent (Ambion, Austin, TX). cDNA was synthesized by reverse transcription using the EasyScript® One-Step gDNA Removal and cDNA Synthesis SuperMix kit (Trans, AE311, China) with oligo dT primer. The cDNA served as a template for real-time qPCR using SYBR green master mix (Dongsheng Biotech, China) and corresponding primers. Real-time qPCR was conducted in the CFX-96 Bio-rad instrument (Bio-rad, USA), and the specificity of the amplified PCR products was confirmed by melting curve analysis after each reaction. The primers used for IFNβ, IFITM3, and IL-8 in this study are listed in Table 3. Statistical analysis was performed using Graphpad Prism8 software. The data are presented as mean ± standard deviation (SD) of three independent experiments. Significance was determined using the two-tailed independent Student’s t-test (P < 0.05) between two groups. One-way analysis of variance followed by Tukey’s test was used to compare multiple groups (>2).

**Table 3.**
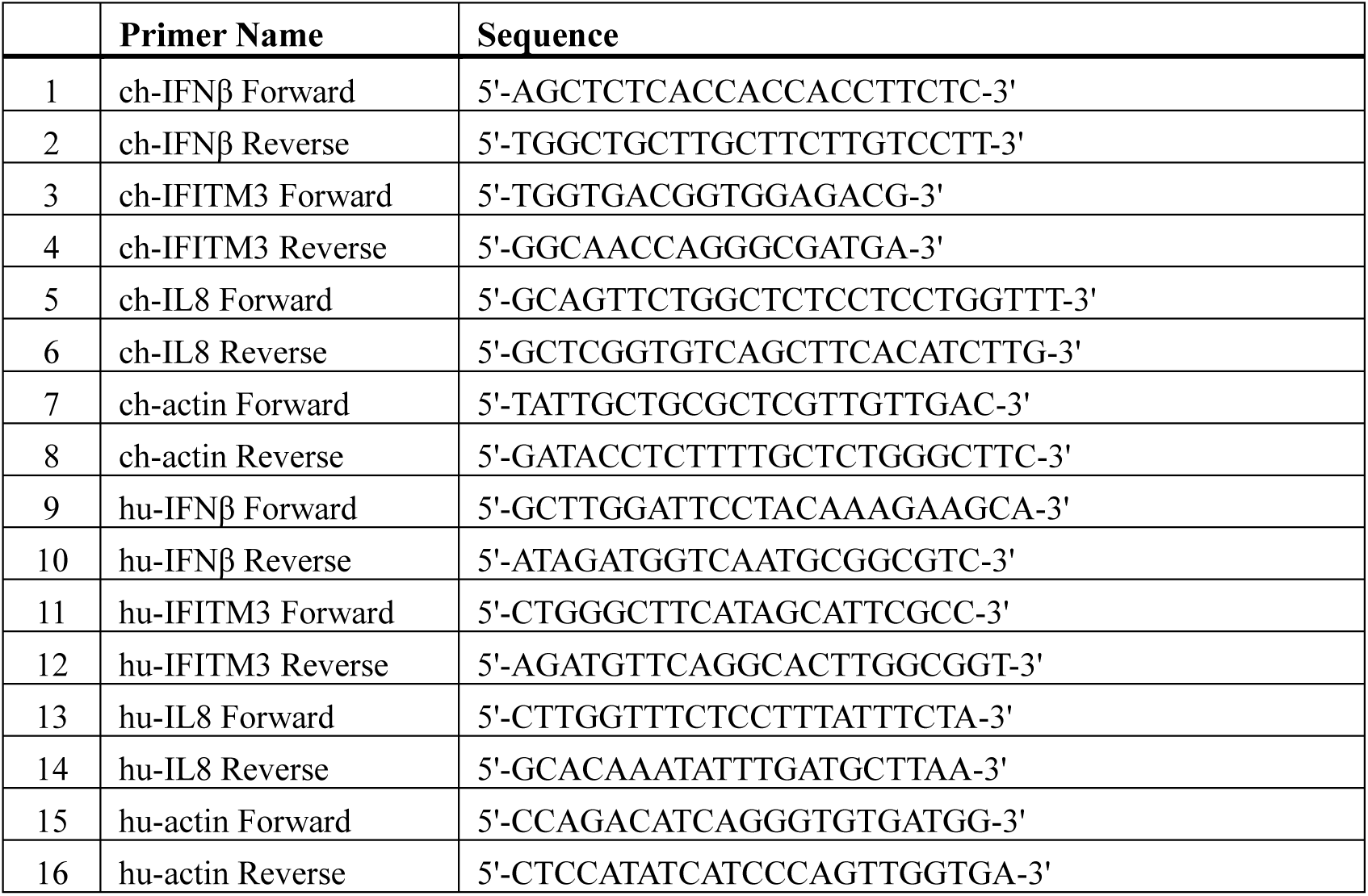
Primer sequences used for real-time qPCR.

### Co-immunoprecipitation (Co-IP) and liquid chromatography-mass spectrometry

DF-1 cells or HEK-293T cells cultured in 6 cm plates were transfected with plasmid or infected with IBV. Cells were lysed using RIPA Lysis Buffer (Beyotime, P0013D, China) supplemented with 1 mM phenylmethylsulfonyl fluoride (PMSF) (Beyotime, ST506, China) and protease inhibitors (Millipore, USA). The cell lysates were centrifuged at 12,000 rpm for 15 min, and the supernatant proteins were incubated overnight at 4°C with gentle rotation with 4 μg of IBV N antibody (mouse), or 2 μg of Flag Tag antibody (mouse) conjugated with Dynabeads Protein G Magnetic Beads (Invitrogen, 10004D, USA), or anti-HA magnetic Beads (Abmart, M20034, China). After incubation, the beads were washed three times with RIPA buffer and then precipitated using a magnetic stand. The beads were resuspended in 40 μL of RIPA lysis buffer and denatured by boiling at 100°C for 5 min after adding 5 × SDS loading buffer (Beyotime, P0015L, China). Following centrifugation, the supernatants were subjected to Western blot analysis.

For the samples intended for Co-IP and liquid chromatography-mass spectrometry (LC-MS) analysis, HEK-293T cells were seeded in 10 cm plates and transfected with Flag-tagged IBV N or PXJ40. At 30 h post-transfection, cells were lysed, and Co-IP experiments were conducted using anti-Flag antibody. Coomassie blue-stained gels from the Co-IP experiments were pooled and subjected to protein identification via LC-MS analysis performed at Jingjie PTM BioLab (Hangzhou, China).

## Acknowledgments

We gratefully acknowledge Prof. Dingxiang Liu from South China Agricultural University, China, for providing the IBV Beaudette strain. Our sincere appreciation also goes to Prof. Tongling Shan from the Shanghai Academy of Agricultural Sciences, CAAS, for generously sharing the plasmids encoding N protein from various genera of coronaviruses. Furthermore, we would like to thank Dr. Huan Wang, also affiliated with the Shanghai Academy of Agricultural Sciences, CAAS, for his contribution in generating the polyclonal IBV N antibody.

## Author Contributions

Conceptualization: Wenxiang Xue, Ying Liao.

Formal analysis: Wenxiang Xue, Ying Liao.

Funding acquisition: Chan Ding, Ying Liao.

Investigation: Wenxiang Xue, Hongyan Chu, Jiehuang Wang.

Project administration: Chan Ding, Ying Liao.

Resources: Yingjie Sun, Lei Tan, Cuiping Song, Xusheng Qiu.

Supervision: Yingjie Sun, Chan Ding, Ying Liao.

Writing-original draft: Wenxiang Xue, Ying Liao.

Writing-review & editing: Ying Liao.

## References

1. Yarbrough ML, Mata MA, Sakthivel R, Fontoura BM. Viral subversion of nucleocytoplasmic trafficking. Traffic. 2014;15(2):127–40. Epub 2013/12/03. doi: 10.1111/tra.12137. PubMed PMID: 24289861; PubMed Central PMCID: PMCPMC3910510.

2. Makhnevych T, Lusk CP, Anderson AM, Aitchison JD, Wozniak RW. Cell cycle regulated transport controlled by alterations in the nuclear pore complex. Cell. 2003;115(7):813–23. Epub 2003/12/31. doi: 10.1016/s0092-8674(03)00986-3. PubMed PMID: 14697200.

3. Sun J, Shi Y, Yildirim E. The Nuclear Pore Complex in Cell Type-Specific Chromatin Structure and Gene Regulation. Trends Genet. 2019;35(8):579–88. Epub 2019/06/20. doi: 10.1016/j.tig.2019.05.006. PubMed PMID: 31213386.

4. Gu Y, Zebell SG, Liang Z, Wang S, Kang BH, Dong X. Nuclear Pore Permeabilization Is a Convergent Signaling Event in Effector-Triggered Immunity. Cell. 2016;166(6):1526–38 e11. Epub 2016/08/30. doi: 10.1016/j.cell.2016.07.042. PubMed PMID: 27569911; PubMed Central PMCID: PMCPMC5017918.

5. Buendia B, Courvalin JC, Collas P. Dynamics of the nuclear envelope at mitosis and during apoptosis. Cell Mol Life Sci. 2001;58(12-13):1781–9. Epub 2002/01/05. doi: 10.1007/PL00000818. PubMed PMID: 11766879.

6. Hampoelz B, Baumbach J. Nuclear envelope assembly and dynamics during development. Semin Cell Dev Biol. 2023;133:96–106. Epub 2022/03/08. doi: 10.1016/j.semcdb.2022.02.028. PubMed PMID: 35249812.

7. Lin DH, Stuwe T, Schilbach S, Rundlet EJ, Perriches T, Mobbs G, et al. Architecture of the symmetric core of the nuclear pore. Science. 2016;352(6283):aaf1015. Epub 2016/04/16. doi: 10.1126/science.aaf1015. PubMed PMID: 27081075; PubMed Central PMCID: PMCPMC5207208.

8. Lin DH, Hoelz A. The Structure of the Nuclear Pore Complex (An Update). Annu Rev Biochem. 2019;88:725–83. Epub 2019/03/19. doi: 10.1146/annurev-biochem-062917-011901. PubMed PMID: 30883195; PubMed Central PMCID: PMCPMC6588426.

9. Beck M, Hurt E. The nuclear pore complex: understanding its function through structural insight. Nat Rev Mol Cell Biol. 2017;18(2):73–89. Epub 2016/12/22. doi: 10.1038/nrm.2016.147. PubMed PMID: 27999437.

10. Shen Q, Wang YE, Palazzo AF. Crosstalk between nucleocytoplasmic trafficking and the innate immune response to viral infection. J Biol Chem. 2021;297(1):100856. Epub 2021/06/08. doi: 10.1016/j.jbc.2021.100856. PubMed PMID: 34097873; PubMed Central PMCID: PMCPMC8254040.

11. Thompson LJ, Fields AP. betaII protein kinase C is required for the G2/M phase transition of cell cycle. J Biol Chem. 1996;271(25):15045–53. Epub 1996/06/21. doi: 10.1074/jbc.271.25.15045. PubMed PMID: 8663071.

12. Collas P. Sequential PKC- and Cdc2-mediated phosphorylation events elicit zebrafish nuclear envelope disassembly. J Cell Sci. 1999;112 ( Pt 6):977–87. Epub 1999/02/26. doi: 10.1242/jcs.112.6.977. PubMed PMID: 10036247.

13. Linder MI, Kohler M, Boersema P, Weberruss M, Wandke C, Marino J, et al. Mitotic Disassembly of Nuclear Pore Complexes Involves CDK1- and PLK1-Mediated Phosphorylation of Key Interconnecting Nucleoporins. Dev Cell. 2017;43(2):141–56 e7. Epub 2017/10/25. doi: 10.1016/j.devcel.2017.08.020. PubMed PMID: 29065306; PubMed Central PMCID: PMCPMC5654724.

14. Martino L, Morchoisne-Bolhy S, Cheerambathur DK, Van Hove L, Dumont J, Joly N, et al. Channel Nucleoporins Recruit PLK-1 to Nuclear Pore Complexes to Direct Nuclear Envelope Breakdown in C. elegans. Dev Cell. 2017;43(2):157–71 e7. Epub 2017/10/25. doi: 10.1016/j.devcel.2017.09.019. PubMed PMID: 29065307; PubMed Central PMCID: PMCPMC8184135.

15. Favreau C, Worman HJ, Wozniak RW, Frappier T, Courvalin JC. Cell cycle-dependent phosphorylation of nucleoporins and nuclear pore membrane protein Gp210. Biochemistry. 1996;35(24):8035–44. Epub 1996/06/18. doi: 10.1021/bi9600660. PubMed PMID: 8672508.

16. Wurzenberger C, Gerlich DW. Phosphatases: providing safe passage through mitotic exit. Nat Rev Mol Cell Biol. 2011;12(8):469–82. Epub 2011/07/14. doi: 10.1038/nrm3149. PubMed PMID: 21750572.

17. Grallert A, Boke E, Hagting A, Hodgson B, Connolly Y, Griffiths JR, et al. A PP1-PP2A phosphatase relay controls mitotic progression. Nature. 2015;517(7532):94-8. Epub 2014/12/10. doi: 10.1038/nature14019. PubMed PMID: 25487150; PubMed Central PMCID: PMCPMC4338534.

18. Xue W, Ding C, Qian K, Liao Y. The Interplay Between Coronavirus and Type I IFN Response. Front Microbiol. 2021;12:805472. Epub 2022/03/24. doi: 10.3389/fmicb.2021.805472. PubMed PMID: 35317429; PubMed Central PMCID: PMCPMC8934427.

19. Schneider WM, Chevillotte MD, Rice CM. Interferon-stimulated genes: a complex web of host defenses. Annu Rev Immunol. 2014;32:513–45. Epub 2014/02/22. doi: 10.1146/annurev-immunol-032713-120231. PubMed PMID: 24555472; PubMed Central PMCID: PMCPMC4313732.

20. Lawrence T. The nuclear factor NF-kappaB pathway in inflammation. Cold Spring Harb Perspect Biol. 2009;1(6):a001651. Epub 2010/05/12. doi: 10.1101/cshperspect.a001651. PubMed PMID: 20457564; PubMed Central PMCID: PMCPMC2882124.

21. Maik-Rachline G, Lifshits L, Seger R. Nuclear P38: Roles in Physiological and Pathological Processes and Regulation of Nuclear Translocation. Int J Mol Sci. 2020;21(17). Epub 2020/08/28. doi: 10.3390/ijms21176102. PubMed PMID: 32847129; PubMed Central PMCID: PMCPMC7504396.

22. Maik-Rachline G, Zehorai E, Hanoch T, Blenis J, Seger R. The nuclear translocation of the kinases p38 and JNK promotes inflammation-induced cancer. Sci Signal. 2018;11(525). Epub 2018/04/11. doi: 10.1126/scisignal.aao3428. PubMed PMID: 29636389.

23. Wang Y, Grunewald M, Perlman S. Coronaviruses: An Updated Overview of Their Replication and Pathogenesis. Methods Mol Biol. 2020;2203:1–29. Epub 2020/08/25. doi: 10.1007/978-1-0716-0900-2_1. PubMed PMID: 32833200; PubMed Central PMCID: PMCPMC7682345.

24. Hasoksuz M, Kilic S, Sarac F. Coronaviruses and SARS-COV-2. Turk J Med Sci. 2020;50(SI-1):549-56. Epub 2020/04/16. doi: 10.3906/sag-2004-127. PubMed PMID: 32293832; PubMed Central PMCID: PMCPMC7195990.

25. Hudson CB, Beaudette FR. Infection of the Cloaca with the Virus of Infectious Bronchitis. Science. 1932;76(1958):34. Epub 1932/07/08. doi: 10.1126/science.76.1958.34-a. PubMed PMID: 17732084.

26. Cavanagh D. Coronavirus avian infectious bronchitis virus. Vet Res. 2007;38(2):281–97. Epub 2007/02/14. doi: 10.1051/vetres:2006055. PubMed PMID: 17296157.

27. Jordan B. Vaccination against infectious bronchitis virus: A continuous challenge. Vet Microbiol. 2017;206:137–43. Epub 2017/01/14. doi: 10.1016/j.vetmic.2017.01.002. PubMed PMID: 28081857.

28. Pensaert MB, de Bouck P. A new coronavirus-like particle associated with diarrhea in swine. Arch Virol. 1978;58(3):243–7. Epub 1978/01/01. doi: 10.1007/BF01317606. PubMed PMID: 83132; PubMed Central PMCID: PMCPMC7086830.

29. Zhang H, Zou C, Peng O, Ashraf U, Xu Q, Gong L, et al. Global Dynamics of Porcine Enteric Coronavirus PEDV Epidemiology, Evolution, and Transmission. Mol Biol Evol. 2023;40(3). Epub 2023/03/05. doi: 10.1093/molbev/msad052. PubMed PMID: 36869744; PubMed Central PMCID: PMCPMC10027654.

30. Jung K, Annamalai T, Lu Z, Saif LJ. Comparative pathogenesis of US porcine epidemic diarrhea virus (PEDV) strain PC21A in conventional 9-day-old nursing piglets vs. 26-day-old weaned pigs. Vet Microbiol. 2015;178(1-2):31–40. Epub 2015/05/06. doi: 10.1016/j.vetmic.2015.04.022. PubMed PMID: 25939885; PubMed Central PMCID: PMCPMC7117181.

31. Cao L, Ge X, Gao Y, Herrler G, Ren Y, Ren X, et al. Porcine epidemic diarrhea virus inhibits dsRNA-induced interferon-beta production in porcine intestinal epithelial cells by blockade of the RIG-I-mediated pathway. Virol J. 2015;12:127. Epub 2015/08/19. doi: 10.1186/s12985-015-0345-x. PubMed PMID: 26283628; PubMed Central PMCID: PMCPMC4539884.

32. Channappanavar R, Fehr AR, Vijay R, Mack M, Zhao J, Meyerholz DK, et al. Dysregulated Type I Interferon and Inflammatory Monocyte-Macrophage Responses Cause Lethal Pneumonia in SARS-CoV-Infected Mice. Cell Host Microbe. 2016;19(2):181–93. Epub 2016/02/13. doi: 10.1016/j.chom.2016.01.007. PubMed PMID: 26867177; PubMed Central PMCID: PMCPMC4752723.

33. Xia H, Cao Z, Xie X, Zhang X, Chen JY, Wang H, et al. Evasion of Type I Interferon by SARS-CoV-2. Cell Rep. 2020;33(1):108234. Epub 2020/09/28. doi: 10.1016/j.celrep.2020.108234. PubMed PMID: 32979938; PubMed Central PMCID: PMCPMC7501843.

34. Lei X, Dong X, Ma R, Wang W, Xiao X, Tian Z, et al. Activation and evasion of type I interferon responses by SARS-CoV-2. Nat Commun. 2020;11(1):3810. Epub 2020/08/01. doi: 10.1038/s41467-020-17665-9. PubMed PMID: 32733001; PubMed Central PMCID: PMCPMC7392898.

35. Channappanavar R, Fehr AR, Zheng J, Wohlford-Lenane C, Abrahante JE, Mack M, et al. IFN-I response timing relative to virus replication determines MERS coronavirus infection outcomes. J Clin Invest. 2019;129(9):3625–39. Epub 2019/07/30. doi: 10.1172/JCI126363. PubMed PMID: 31355779; PubMed Central PMCID: PMCPMC6715373.

36. Roth-Cross JK, Martinez-Sobrido L, Scott EP, Garcia-Sastre A, Weiss SR. Inhibition of the alpha/beta interferon response by mouse hepatitis virus at multiple levels. J Virol. 2007;81(13):7189–99. Epub 2007/04/27. doi: 10.1128/JVI.00013-07. PubMed PMID: 17459917; PubMed Central PMCID: PMCPMC1933268.

37. Kint J, Fernandez-Gutierrez M, Maier HJ, Britton P, Langereis MA, Koumans J, et al. Activation of the chicken type I interferon response by infectious bronchitis coronavirus. J Virol. 2015;89(2):1156–67. Epub 2014/11/08. doi: 10.1128/JVI.02671-14. PubMed PMID: 25378498; PubMed Central PMCID: PMCPMC4300645.

38. Luo J, Fang L, Dong N, Fang P, Ding Z, Wang D, et al. Porcine deltacoronavirus (PDCoV) infection suppresses RIG-I-mediated interferon-beta production. Virology. 2016;495:10–7. Epub 2016/05/07. doi: 10.1016/j.virol.2016.04.025. PubMed PMID: 27152478; PubMed Central PMCID: PMCPMC7111668.

39. Zhang K, Lin S, Li J, Deng S, Zhang J, Wang S. Modulation of Innate Antiviral Immune Response by Porcine Enteric Coronavirus. Front Microbiol. 2022;13:845137. Epub 2022/03/04. doi: 10.3389/fmicb.2022.845137. PubMed PMID: 35237253; PubMed Central PMCID: PMCPMC8882816.

40. Lowery SA, Sariol A, Perlman S. Innate immune and inflammatory responses to SARS-CoV-2: Implications for COVID-19. Cell Host Microbe. 2021;29(7):1052–62. Epub 2021/05/23. doi: 10.1016/j.chom.2021.05.004. PubMed PMID: 34022154; PubMed Central PMCID: PMCPMC8126603.

41. Frieman M, Heise M, Baric R. SARS coronavirus and innate immunity. Virus Res. 2008;133(1):101-12. Epub 2007/04/25. doi: 10.1016/j.virusres.2007.03.015. PubMed PMID: 17451827; PubMed Central PMCID: PMCPMC2292640.

42. Frieman M, Yount B, Heise M, Kopecky-Bromberg SA, Palese P, Baric RS. Severe acute respiratory syndrome coronavirus ORF6 antagonizes STAT1 function by sequestering nuclear import factors on the rough endoplasmic reticulum/Golgi membrane. J Virol. 2007;81(18):9812–24. Epub 2007/06/29. doi: 10.1128/JVI.01012-07. PubMed PMID: 17596301; PubMed Central PMCID: PMCPMC2045396.

43. Miorin L, Kehrer T, Sanchez-Aparicio MT, Zhang K, Cohen P, Patel RS, et al. SARS-CoV-2 Orf6 hijacks Nup98 to block STAT nuclear import and antagonize interferon signaling. Proc Natl Acad Sci U S A. 2020;117(45):28344–54. Epub 2020/10/25. doi: 10.1073/pnas.2016650117. PubMed PMID: 33097660; PubMed Central PMCID: PMCPMC7668094.

44. Kint J, Dickhout A, Kutter J, Maier HJ, Britton P, Koumans J, et al. Infectious Bronchitis Coronavirus Inhibits STAT1 Signaling and Requires Accessory Proteins for Resistance to Type I Interferon Activity. J Virol. 2015;89(23):12047–57. Epub 2015/09/25. doi: 10.1128/JVI.01057-15. PubMed PMID: 26401035; PubMed Central PMCID: PMCPMC4645315.

45. Emeny JM, Morgan MJ. Regulation of the interferon system: evidence that Vero cells have a genetic defect in interferon production. J Gen Virol. 1979;43(1):247–52. Epub 1979/04/01. doi: 10.1099/0022-1317-43-1-247. PubMed PMID: 113494.

46. Mosca JD, Pitha PM. Transcriptional and posttranscriptional regulation of exogenous human beta interferon gene in simian cells defective in interferon synthesis. Mol Cell Biol. 1986;6(6):2279–83. Epub 1986/06/01. doi: 10.1128/mcb.6.6.2279-2283.1986. PubMed PMID: 3785197; PubMed Central PMCID: PMCPMC367773.

47. De Jesus-Gonzalez LA, Cervantes-Salazar M, Reyes-Ruiz JM, Osuna-Ramos JF, Farfan-Morales CN, Palacios-Rapalo SN, et al. The Nuclear Pore Complex: A Target for NS3 Protease of Dengue and Zika Viruses. Viruses. 2020;12(6). Epub 2020/05/30. doi: 10.3390/v12060583. PubMed PMID: 32466480; PubMed Central PMCID: PMCPMC7354628.

48. Kutay U, Juhlen R, Antonin W. Mitotic disassembly and reassembly of nuclear pore complexes. Trends Cell Biol. 2021;31(12):1019–33. Epub 2021/07/24. doi: 10.1016/j.tcb.2021.06.011. PubMed PMID: 34294532.

49. Malik YA. Properties of Coronavirus and SARS-CoV-2. Malays J Pathol. 2020;42(1):3–11. Epub 2020/04/29. PubMed PMID: 32342926.

50. Emmott E, Munday D, Bickerton E, Britton P, Rodgers MA, Whitehouse A, et al. The cellular interactome of the coronavirus infectious bronchitis virus nucleocapsid protein and functional implications for virus biology. J Virol. 2013;87(17):9486–500. Epub 2013/05/03. doi: 10.1128/JVI.00321-13. PubMed PMID: 23637410; PubMed Central PMCID: PMCPMC3754094.

51. Zhou J, Qiu Y, Zhao J, Wang Y, Zhu N, Wang D, et al. The Network of Interactions between the Porcine Epidemic Diarrhea Virus Nucleocapsid and Host Cellular Proteins. Viruses. 2022;14(10). Epub 2022/10/28. doi: 10.3390/v14102269. PubMed PMID: 36298827; PubMed Central PMCID: PMCPMC9611260.

52. Zheng X, Sun Z, Yu L, Shi D, Zhu M, Yao H, et al. Interactome Analysis of the Nucleocapsid Protein of SARS-CoV-2 Virus. Pathogens. 2021;10(9). Epub 2021/09/29. doi: 10.3390/pathogens10091155. PubMed PMID: 34578187; PubMed Central PMCID: PMCPMC8465953.

53. Min YQ, Huang M, Feng K, Jia Y, Sun X, Ning YJ. A New Cellular Interactome of SARS-CoV-2 Nucleocapsid Protein and Its Biological Implications. Mol Cell Proteomics. 2023;22(7):100579. Epub 2023/05/22. doi: 10.1016/j.mcpro.2023.100579. PubMed PMID: 37211047; PubMed Central PMCID: PMCPMC10198743.

54. Adams DR, Ron D, Kiely PA. RACK1, A multifaceted scaffolding protein: Structure and function. Cell Commun Signal. 2011;9:22. Epub 2011/10/08. doi: 10.1186/1478-811X-9-22. PubMed PMID: 21978545; PubMed Central PMCID: PMCPMC3195729.

55. Hocevar BA, Burns DJ, Fields AP. Identification of protein kinase C (PKC) phosphorylation sites on human lamin B. Potential role of PKC in nuclear lamina structural dynamics. J Biol Chem. 1993;268(10):7545–52. Epub 1993/04/05. PubMed PMID: 8463284.

56. Eggert M, Radomski N, Linder D, Tripier D, Traub P, Jost E. Identification of novel phosphorylation sites in murine A-type lamins. Eur J Biochem. 1993;213(2):659–71. Epub 1993/04/15. doi: 10.1111/j.1432-1033.1993.tb17806.x. PubMed PMID: 8477740.

57. Livneh E, Fishman DD. Linking protein kinase C to cell-cycle control. Eur J Biochem. 1997;248(1):1–9. Epub 1997/08/15. doi: 10.1111/j.1432-1033.1997.t01-4-00001.x. PubMed PMID: 9310352.

58. Newton AC. Protein kinase C: poised to signal. Am J Physiol Endocrinol Metab. 2010;298(3):E395–402. Epub 2009/11/26. doi: 10.1152/ajpendo.00477.2009. PubMed PMID: 19934406; PubMed Central PMCID: PMCPMC2838521.

59. Newton AC. Regulation of protein kinase C. Curr Opin Cell Biol. 1997;9(2):161–7. Epub 1997/04/01. doi: 10.1016/s0955-0674(97)80058-0. PubMed PMID: 9069266.

60. Solmaz SR, Chauhan R, Blobel G, Melcak I. Molecular architecture of the transport channel of the nuclear pore complex. Cell. 2011;147(3):590–602. Epub 2011/11/01. doi: 10.1016/j.cell.2011.09.034. PubMed PMID: 22036567; PubMed Central PMCID: PMCPMC3431207.

61. Stuwe T, Bley CJ, Thierbach K, Petrovic S, Schilbach S, Mayo DJ, et al. Architecture of the fungal nuclear pore inner ring complex. Science. 2015;350(6256):56-64. Epub 2015/09/01. doi: 10.1126/science.aac9176. PubMed PMID: 26316600; PubMed Central PMCID: PMCPMC4826903.

62. Schechtman D, Mochly-Rosen D. Adaptor proteins in protein kinase C-mediated signal transduction. Oncogene. 2001;20(44):6339–47. Epub 2001/10/19. doi: 10.1038/sj.onc.1204778. PubMed PMID: 11607837.

63. Hampoelz B, Andres-Pons A, Kastritis P, Beck M. Structure and Assembly of the Nuclear Pore Complex. Annu Rev Biophys. 2019;48:515–36. Epub 2019/04/04. doi: 10.1146/annurev-biophys-052118-115308. PubMed PMID: 30943044.

64. Hayama R, Rout MP, Fernandez-Martinez J. The nuclear pore complex core scaffold and permeability barrier: variations of a common theme. Curr Opin Cell Biol. 2017;46:110–8. Epub 2017/06/19. doi: 10.1016/j.ceb.2017.05.003. PubMed PMID: 28624666; PubMed Central PMCID: PMCPMC5568245.

65. Porter FW, Palmenberg AC. Leader-induced phosphorylation of nucleoporins correlates with nuclear trafficking inhibition by cardioviruses. J Virol. 2009;83(4):1941–51. Epub 2008/12/17. doi: 10.1128/JVI.01752-08. PubMed PMID: 19073724; PubMed Central PMCID: PMCPMC2643766.

66. Porter FW, Brown B, Palmenberg AC. Nucleoporin phosphorylation triggered by the encephalomyocarditis virus leader protein is mediated by mitogen-activated protein kinases. J Virol. 2010;84(24):12538–48. Epub 2010/10/01. doi: 10.1128/JVI.01484-09. PubMed PMID: 20881039; PubMed Central PMCID: PMCPMC3004318.

67. Lu D, Yang H, Raizada MK. Involvement of p62 nucleoporin in angiotensin II-induced nuclear translocation of STAT3 in brain neurons. J Neurosci. 1998;18(4):1329–36. Epub 1998/03/14. doi: 10.1523/JNEUROSCI.18-04-01329.1998. PubMed PMID: 9454842; PubMed Central PMCID: PMCPMC6792719.

68. Porwal M, Cohen S, Snoussi K, Popa-Wagner R, Anderson F, Dugot-Senant N, et al. Parvoviruses cause nuclear envelope breakdown by activating key enzymes of mitosis. PLoS Pathog. 2013;9(10):e1003671. Epub 2013/11/10. doi: 10.1371/journal.ppat.1003671. PubMed PMID: 24204256; PubMed Central PMCID: PMCPMC3814971.

69. Kawano T, Inokuchi J, Eto M, Murata M, Kang JH. Activators and Inhibitors of Protein Kinase C (PKC): Their Applications in Clinical Trials. Pharmaceutics. 2021;13(11). Epub 2021/11/28. doi: 10.3390/pharmaceutics13111748. PubMed PMID: 34834162; PubMed Central PMCID: PMCPMC8621927.

70. Kawauchi K, Lazarus AH, Sanghera JS, Man GL, Pelech SL, Delovitch TL. Regulation of BCR-and PKC/Ca(2+)-mediated activation of the Raf1/MEK/MAPK pathway by protein-tyrosine kinase and -tyrosine phosphatase activities. Mol Immunol. 1996;33(3):287–96. Epub 1996/02/01. doi: 10.1016/0161-5890(95)00134-4. PubMed PMID: 8649450.

71. Jiffar T, Kurinna S, Suck G, Carlson-Bremer D, Ricciardi MR, Konopleva M, et al. PKC alpha mediates chemoresistance in acute lymphoblastic leukemia through effects on Bcl2 phosphorylation. Leukemia. 2004;18(3):505–12. Epub 2004/01/23. doi: 10.1038/sj.leu.2403275. PubMed PMID: 14737078.

72. Xu X, Han L, Zhao G, Xue S, Gao Y, Xiao J, et al. LRCH1 interferes with DOCK8-Cdc42-induced T cell migration and ameliorates experimental autoimmune encephalomyelitis. J Exp Med. 2017;214(1):209–26. Epub 2016/12/29. doi: 10.1084/jem.20160068. PubMed PMID: 28028151; PubMed Central PMCID: PMCPMC5206493.

73. Albano GD, Bonanno A, Moscato M, Anzalone G, Di Sano C, Riccobono L, et al. Crosstalk between mAChRM3 and beta2AR, via acetylcholine PI3/PKC/PBEP1/Raf-1 MEK1/2/ERK1/2 pathway activation, in human bronchial epithelial cells after long-term cigarette smoke exposure. Life Sci. 2018;192:99–109. Epub 2017/11/28. doi: 10.1016/j.lfs.2017.11.034. PubMed PMID: 29175450.

74. Ghandour H, Cullere X, Alvarez A, Luscinskas FW, Mayadas TN. Essential role for Rap1 GTPase and its guanine exchange factor CalDAG-GEFI in LFA-1 but not VLA-4 integrin mediated human T-cell adhesion. Blood. 2007;110(10):3682–90. Epub 2007/08/19. doi: 10.1182/blood-2007-03-077628. PubMed PMID: 17702895; PubMed Central PMCID: PMCPMC2077316.

75. Huang C, Feng F, Shi Y, Li W, Wang Z, Zhu Y, et al. Protein Kinase C Inhibitors Reduce SARS-CoV-2 Replication in Cultured Cells. Microbiol Spectr. 2022;10(5):e0105622. Epub 2022/08/25. doi: 10.1128/spectrum.01056-22. PubMed PMID: 36000889; PubMed Central PMCID: PMCPMC9603170.

76. Ron D, Luo J, Mochly-Rosen D. C2 region-derived peptides inhibit translocation and function of beta protein kinase C in vivo. J Biol Chem. 1995;270(41):24180–7. Epub 1995/10/13. doi: 10.1074/jbc.270.41.24180. PubMed PMID: 7592622.

77. Shue B, Chiramel AI, Cerikan B, To TH, Frolich S, Pederson SM, et al. Genome-Wide CRISPR Screen Identifies RACK1 as a Critical Host Factor for Flavivirus Replication. J Virol. 2021;95(24):e0059621. Epub 2021/09/30. doi: 10.1128/JVI.00596-21. PubMed PMID: 34586867; PubMed Central PMCID: PMCPMC8610583.

78. Wang X, Bi J, Yang Y, Li L, Zhang R, Li Y, et al. RACK1 promotes porcine reproductive and respiratory syndrome virus infection in Marc-145 cells through ERK1/2 activation. Virology. 2023;588:109886. Epub 2023/10/09. doi: 10.1016/j.virol.2023.109886. PubMed PMID: 37806007.

79. Stebbins EG, Mochly-Rosen D. Binding specificity for RACK1 resides in the V5 region of beta II protein kinase C. J Biol Chem. 2001;276(32):29644–50. Epub 2001/06/02. doi: 10.1074/jbc.M101044200. PubMed PMID: 11387319.

80. McBride R, van Zyl M, Fielding BC. The coronavirus nucleocapsid is a multifunctional protein. Viruses. 2014;6(8):2991–3018. Epub 2014/08/12. doi: 10.3390/v6082991. PubMed PMID: 25105276; PubMed Central PMCID: PMCPMC4147684.

81. Bai Z, Cao Y, Liu W, Li J. The SARS-CoV-2 Nucleocapsid Protein and Its Role in Viral Structure, Biological Functions, and a Potential Target for Drug or Vaccine Mitigation. Viruses. 2021;13(6). Epub 2021/07/03. doi: 10.3390/v13061115. PubMed PMID: 34200602; PubMed Central PMCID: PMCPMC8227405.

82. Masters PS. Localization of an RNA-binding domain in the nucleocapsid protein of the coronavirus mouse hepatitis virus. Arch Virol. 1992;125(1-4):141–60. Epub 1992/01/01. doi: 10.1007/BF01309634. PubMed PMID: 1322650; PubMed Central PMCID: PMCPMC7086615.

83. Hartenian E, Nandakumar D, Lari A, Ly M, Tucker JM, Glaunsinger BA. The molecular virology of coronaviruses. J Biol Chem. 2020;295(37):12910–34. Epub 2020/07/15. doi: 10.1074/jbc.REV120.013930. PubMed PMID: 32661197; PubMed Central PMCID: PMCPMC7489918.

84. Huang Q, Yu L, Petros AM, Gunasekera A, Liu Z, Xu N, et al. Structure of the N-terminal RNA-binding domain of the SARS CoV nucleocapsid protein. Biochemistry. 2004;43(20):6059–63. Epub 2004/05/19. doi: 10.1021/bi036155b. PubMed PMID: 15147189.

85. Wurm T, Chen H, Hodgson T, Britton P, Brooks G, Hiscox JA. Localization to the nucleolus is a common feature of coronavirus nucleoproteins, and the protein may disrupt host cell division. J Virol. 2001;75(19):9345–56. Epub 2001/09/05. doi: 10.1128/JVI.75.19.9345-9356.2001. PubMed PMID: 11533198; PubMed Central PMCID: PMCPMC114503.

86. Ding Z, Luo S, Gong W, Wang L, Ding N, Chen J, et al. Subcellular localization of the porcine deltacoronavirus nucleocapsid protein. Virus Genes. 2020;56(6):687–95. Epub 2020/09/19. doi: 10.1007/s11262-020-01790-0. PubMed PMID: 32944812; PubMed Central PMCID: PMCPMC7497858.

87. Surjit M, Kumar R, Mishra RN, Reddy MK, Chow VT, Lal SK. The severe acute respiratory syndrome coronavirus nucleocapsid protein is phosphorylated and localizes in the cytoplasm by 14-3-3-mediated translocation. J Virol. 2005;79(17):11476–86. Epub 2005/08/17. doi: 10.1128/JVI.79.17.11476-11486.2005. PubMed PMID: 16103198; PubMed Central PMCID: PMCPMC1193639.

88. Hall R, Guedan A, Yap MW, Young GR, Harvey R, Stoye JP, et al. SARS-CoV-2 ORF6 disrupts innate immune signalling by inhibiting cellular mRNA export. PLoS Pathog. 2022;18(8):e1010349. Epub 2022/08/26. doi: 10.1371/journal.ppat.1010349. PubMed PMID: 36007063; PubMed Central PMCID: PMCPMC9451085.

89. Jiao Y, Kong N, Wang H, Sun D, Dong S, Chen X, et al. PABPC4 Broadly Inhibits Coronavirus Replication by Degrading Nucleocapsid Protein through Selective Autophagy. Microbiol Spectr. 2021;9(2):e0090821. Epub 2021/10/07. doi: 10.1128/Spectrum.00908-21. PubMed PMID: 34612687; PubMed Central PMCID: PMCPMC8510267.

90. Gao B, Gong X, Fang S, Weng W, Wang H, Chu H, et al. Inhibition of anti-viral stress granule formation by coronavirus endoribonuclease nsp15 ensures efficient virus replication. PLoS Pathog. 2021;17(2):e1008690. Epub 2021/02/27. doi: 10.1371/journal.ppat.1008690. PubMed PMID: 33635931; PubMed Central PMCID: PMCPMC7946191.

